# Shared features underlying compact genomes and extreme habitat use in chironomid midges

**DOI:** 10.1101/2023.12.12.571347

**Authors:** Lucas A. Nell, Yi-Ming Weng, Joseph S. Phillips, Jamieson C. Botsch, K. Riley Book, Árni Einarsson, Anthony R. Ives, Sean D. Schoville

**Author notes:** Author for Correspondence: Lucas A. Nell, Department of Biology, Stanford University, Stanford, CA 94305, USA.

## Abstract

Non-biting midges (family Chironomidae) are found throughout the world in a diverse array of aquatic and terrestrial habitats, can often tolerate harsh conditions such as hypoxia or desiccation, and have consistently compact genomes. Yet we know little about the shared molecular basis for these attributes and how they have evolved across the family. Here, we address these questions by first creating high-quality, annotated reference assemblies for *Tanytarsus gracilentus* (subfamily Chironominae, tribe Tanytarsini) and *Parochlus steinenii* (subfamily Podonominae). Using these and other publicly available assemblies, we created a time-calibrated phylogenomic tree for family Chironomidae with outgroups from order Diptera. We used this phylogeny to test for features associated with compact genomes, as well as examining patterns of gene family evolution and positive selection that may underlie chironomid habitat tolerances. Our results suggest that compact genomes evolved in the most recent common ancestor of Chironomidae and Ceratopogonidae, and that this occurred mainly through reductions in non-coding regions (introns, intergenic sequences, and repeat elements). Gene families that significantly expanded in Chironomidae included biological processes that may relate to tolerance of stressful environments, such as temperature homeostasis, inflammatory response, melanization defense response, and trehalose transport. We identified a number of genes with evidence for positive selection in Chironomidae, notably sulfonylurea receptor, peroxiredoxin-1, and protein kinase D. Our results help to understand the genomic basis for the small genomes and extreme habitat use in this widely distributed group.

**Significance Statement:** Chironomid midges are known for having small genomes and being able to tolerate many forms of environmental stress, yet little is known of the shared features of their genomes that may underlie these traits. We found that reductions in non-coding regions coincide with small chironomid genomes, and we identified duplicated and/or selected genes that may equip chironomids to tolerate harsh conditions. These results describe the key genomic changes in chironomid midges that may explain their ability to inhabit a range of extreme habitats across the world.

## Introduction

Non-biting midges of the family Chironomidae (order Diptera) are the most widely distributed group of freshwater insects (Armitage et al. 1995), with species found as far north as Ellesmere Island in Canada (Oliver & Corbet 1966); as far south as the Antarctic mainland (Usher & Edwards 1984); at 5,600 meters above sea level in Himalayan glaciers (Kohshima 1984); and at 1,000 meters below the surface of Lake Baikal in Siberia (Linevich 1963). Sediment in a productive freshwater river or lake is the archetypical chironomid habitat, but chironomids have evolved to live in diverse environments such as ephemeral pools, hot springs, and water-filled cavities in plants, with some species even being fully marine or terrestrial (Armitage et al. 1995). Part of why chironomids are so ubiquitous is their ability, as a group, to tolerate extreme conditions. Many forms of tolerance have been documented, including to low temperature (Lee et al. 2006; Rinehart et al. 2006), low oxygen (Burmester & Hankeln 2007), heavy metal contamination (Zheng et al. 2017), complete desiccation (Watanabe et al. 2002), and exposure to ionizing radiation (Gusev et al. 2010).

The extent to which shared genomic features underlie the extreme physiology and diverse habitat use of chironomids is largely unknown. The first sequenced chironomid genome was the Antarctic midge *Belgica antarctica*, one of the smallest recorded insect genomes (Kelley et al. 2014). The number of protein-coding genes in the *B. antarctica* genome was similar to other non-chironomid dipterans, whereas repeat elements and non-coding regions were reduced, and the authors concluded that the small genome size was likely an adaptation to its extreme environment. However, genome sizes were later estimated for 25 chironomid species by flow cytometry, showing that small genome size is likely an ancestral trait in chironomids and that smaller genomes within the family do not necessarily correlate with particularly stressful environments (Cornette et al. 2015). Additional reference assemblies have yielded further insights into individual types of stress tolerance, including haemoglobin gene repeats underlying copper tolerance in *Propsilocerus akamusi* (Sun et al. 2021) or a single chromosome in *Polypedilum vanderplanki* containing clusters of duplicated genes mediating desiccation tolerance (Gusev et al. 2014; Yoshida et al. 2022). Some commonalities have emerged, such as upregulation in antioxidant genes as a common mechanism for chironomid tolerances to heavy metal exposure, desiccation, and cold (Lopez-Martinez et al. 2008; Gusev et al. 2014; Zheng et al. 2011). Nonetheless, no study has explicitly compared chironomids to other dipterans to help understand genome evolution in this ubiquitous group.

Here, we generated two high-quality, annotated chironomid reference assemblies, a new assembly for *Tanytarsus gracilentus* (subfamily Chironominae, tribe Tanytarsini) and an improved assembly for *Parochlus steinenii* (subfamily Podonominae). We also constructed a time-calibrated phylogenomic tree for nine chironomids plus five dipteran outgroups. We used this phylogeny to inform analyses of (1) genome features associated with compact chironomid genomes and (2) gene family evolution and positive selection that relate to extreme chironomid habitat use.

## Results and Discussion

### Genome Assemblies

We generated 23.08 Gb (∼243×) Oxford Nanopore Technologies (ONT) reads (read length N50 = 7,539 bp) and 22.11 Gb (∼232×) Illumina DNAseq reads from our samples of *Tanytarsus gracilentus*. The resulting assembly was 91.83 Mb in size, which closely matches the size estimated by back-mapping reads to the final assembly (95.10 Mb). We also created an assembly for *Parochlus steinenii* based on sequencing reads from the Sequence Read Archive (SRA). This assembly’s size of 143.57 Mb closely matches the previously published prediction of 143.8 Mb (Shin et al. 2019). These assemblies are two of the most contiguous gap-free chironomid assemblies to date (table 1), although they are not arranged onto chromosomes as in *Polypedilum vanderplanki* and *Propsilocerus akamusi* (Sun et al. 2021; Yoshida et al. 2022). Their high BUSCO (diptera_odb10 library) completeness percentages (*T. gracilentus* = 91.60% [90.53% single-copy, 1.07% duplicated], *P. steinenii* = 92.30% [90.99% single-copy, 1.31% duplicated]) also indicate high quality assemblies. Both assemblies had negligible contamination from other species (maximum from sendsketch.sh: *T. gracilentus* = 0.02%, *P. steinenii* = 0.01%).

**Table 1.**
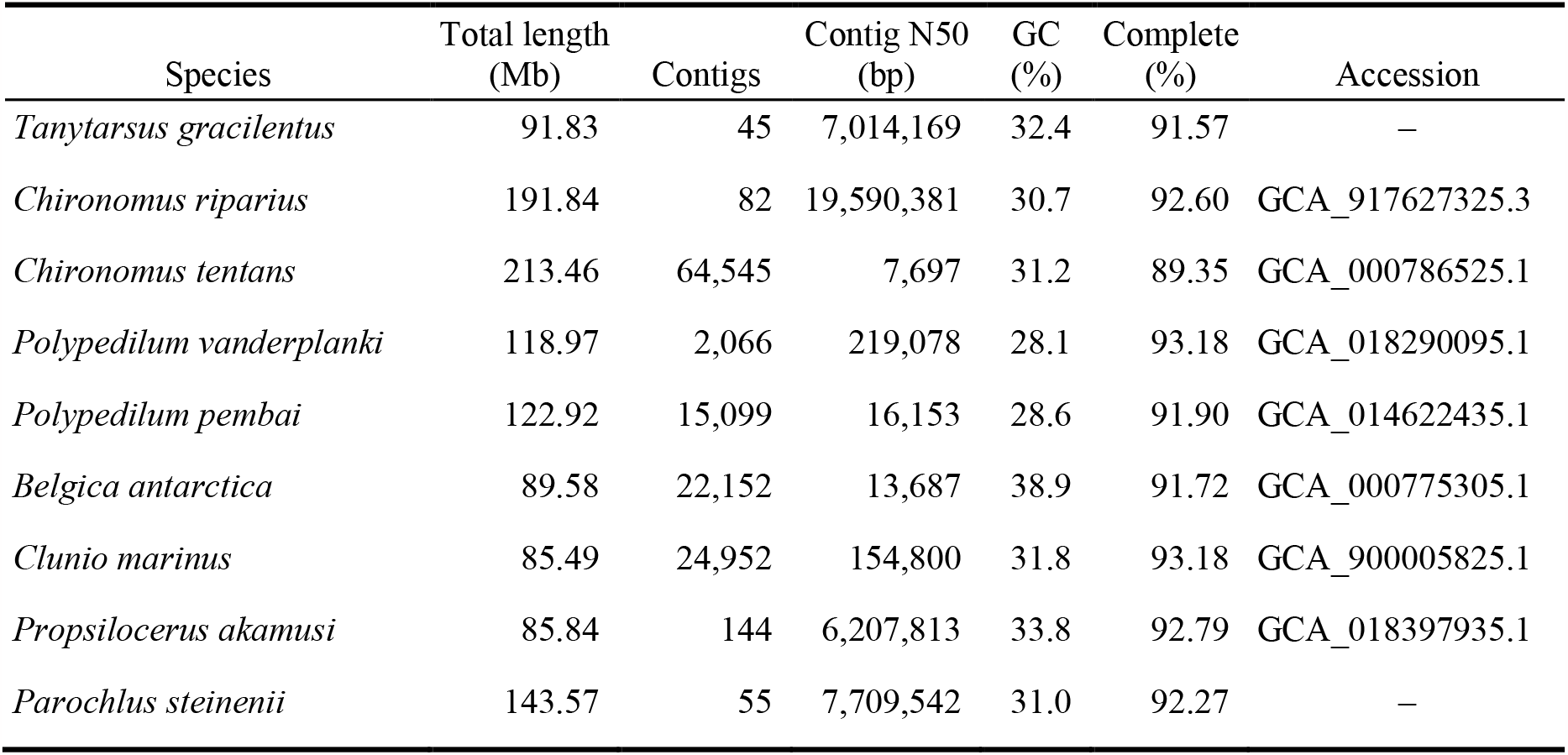
Summary statistics for the final *Tanytarsus gracilentus and Parochlus steinenii* assemblies and for all assemblies of species in the family Chironomidae that are available on GenBank. Percent complete was calculated using BUSCO complete genes based on the diptera_odb10 library.

| Species | Total length (Mb) | Contigs | Contig N50 (bp) | GC (%) | Complete (%) | Accession |
| --- | --- | --- | --- | --- | --- | --- |
| *Tanytarsus gracilentus* | 91.83 | 45 | 7,014,169 | 32.4 | 91.57 | – |
| *Chironomus riparius* | 191.84 | 82 | 19,590,381 | 30.7 | 92.60 | GCA_917627325.3 |
| *Chironomus tentans* | 213.46 | 64,545 | 7,697 | 31.2 | 89.35 | GCA_000786525.1 |
| *Polypedilum vanderplanki* | 118.97 | 2,066 | 219,078 | 28.1 | 93.18 | GCA_018290095.1 |
| *Polypedilum pembai* | 122.92 | 15,099 | 16,153 | 28.6 | 91.90 | GCA_014622435.1 |
| *Belgica antarctica* | 89.58 | 22,152 | 13,687 | 38.9 | 91.72 | GCA_000775305.1 |
| *Clunio marinus* | 85.49 | 24,952 | 154,800 | 31.8 | 93.18 | GCA_900005825.1 |
| *Propiloscerus akamusi* | 85.84 | 144 | 6,207,813 | 33.8 | 92.79 | GCA_018397935.1 |
| *Parochlus steinenii* | 143.57 | 55 | 7,709,542 | 31.0 | 92.27 | – |

### Genome annotations

For *T. gracilentus*, we generated 52.81 Gb (∼555×) Illumina RNAseq reads from adults and 63.28 Gb (∼665×) Illumina RNAseq reads from juveniles. The total protein-coding genes for the gene predictions we created for *T. gracilentus, Parochlus steinenii*, and *Culicoides sonorensis* were similar to other chironomids and dipteran outgroups (table S1). BUSCO completeness was high for the coding sequences from all sets of gene predictions, indicating good performance of the gene predictions, but the number of duplicated genes in *C. sonorensis* was high (fig. S1), likely as a result of its redundant assembly (12.5% BUSCO complete+duplicate genes). We were able to functionally annotate most of the proteins for each species (*C. riparius* = 12,249; *C. sonorensis* = 12,971; *P. steinenii* = 11,645; and *T. gracilentus* = 11,369) with specific tags such as Gene Ontology (GO) terms (table S1).

### Phylogeny

Protein sequences from 1,437 diptera_odb10 single-copy, orthologous genes (1,077,591 total aligned sites after fully undetermined columns removed) were used to construct the 14-species phylogenomic tree of chironomids with dipteran outgroups. All internal nodes were supported with high bootstrap values (fig. S2), and the topology of Chironomidae is consistent with previously published phylogenies (Cranston et al. 2012, 2010). MCMCTree results were consistent across runs, with the minimum pairwise correlation between mean posterior estimates being *r* = 1.000. Our deepest Chironomidae divergence times (fig. 1A) are similar to those in Cranston *et al*. (2012), but more nested splits, such as between *Propsilocerus* and the subfamilies Orthocladiinae and Chironominae, are intermediate when compared to two published estimates (Cranston et al. 2012, 2010). The confidence intervals for our three deepest divergence events were overlapping, which is consistent with other studies showing a wide range of estimates for the divergence times between, for example, Chironomidae and Ceratopogonidae (137.3–296.9 Ma) (Rainford et al. 2014; Bertone et al. 2008; Cranston et al. 2010, 2012).

**Figure 1.**
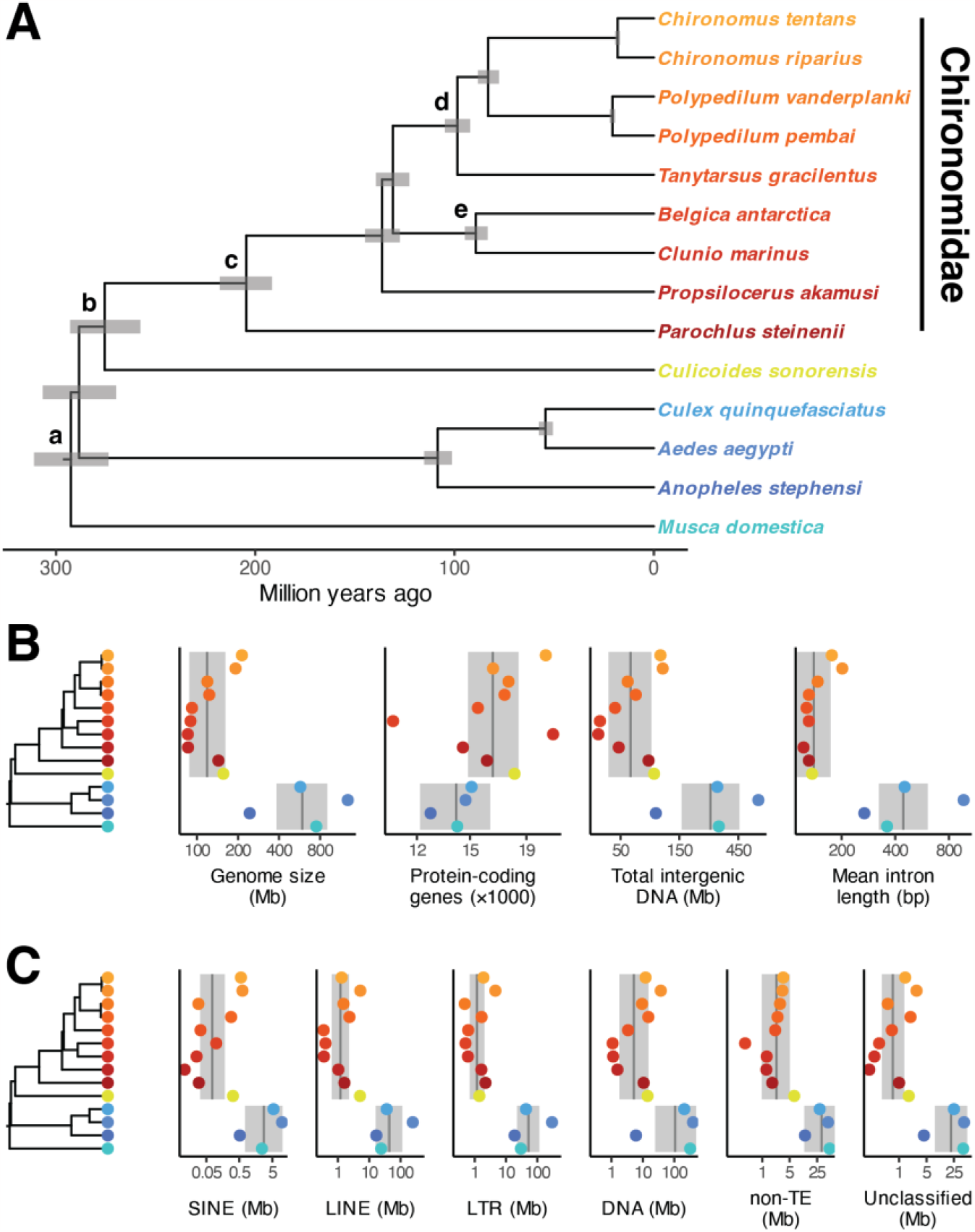
(A) Time-calibrated phylogenomic tree showing relationships among chironomids and outgroups within order Diptera. Gray bars indicate 95% credible intervals for node ages, and lowercase letters indicate fossil calibrations described in the Materials and Methods. Label colors indicate the colors in the lower panels; chironomids are shades of red / orange. (B) Time-calibrated phylogeny next to genome size, number of protein-coding genes, total intergenic content, and mean intron length for all species in our phylogeny. (C) Phylogeny alongside repeat content by class. (B,C) Gray vertical lines indicate the mean estimates from the phylogenetic linear regressions for Chironomidae and Ceratopogonidae and for all other species. Gray envelopes indicate the 95% confidence interval bounds for these estimates computed via parametric bootstrapping. All measures are the log_10_-transformed totals across each species’ entire genome except for intron length, which is the mean of log_10_-transformed intron lengths.

### Features associated with genome size

Genome size, intergenic sequences, introns, and repeat elements all differed between non-chironomid dipterans and the group comprising Chironomidae and their closest relative, *C. sonorensis* (family Ceratopogonidae) (fig. 1B,C). These same features were also significantly correlated with genome size across all species in our phylogeny (fig. 2). In contrast, protein-coding genes neither correlated with genome size nor differed between the families Chironomidae and Ceratopogonidae and other dipterans. These results suggest that the compact genomes of chironomids likely evolved in a common ancestor with Ceratopogonidae, although having only one species from Ceratopogonidae in our analysis makes this conclusion less certain. Our results also suggest that the reduction in genome size occurred through non-coding regions and repeat elements, which is consistent with a recent analysis of contributors to insect genome sizes (Cong et al. 2022). However, unlike Cong et al. (2022), our repeat-element divergence landscapes reveal no obvious connection between repeat element ages and genome size (fig. S3).

**Figure 2.**
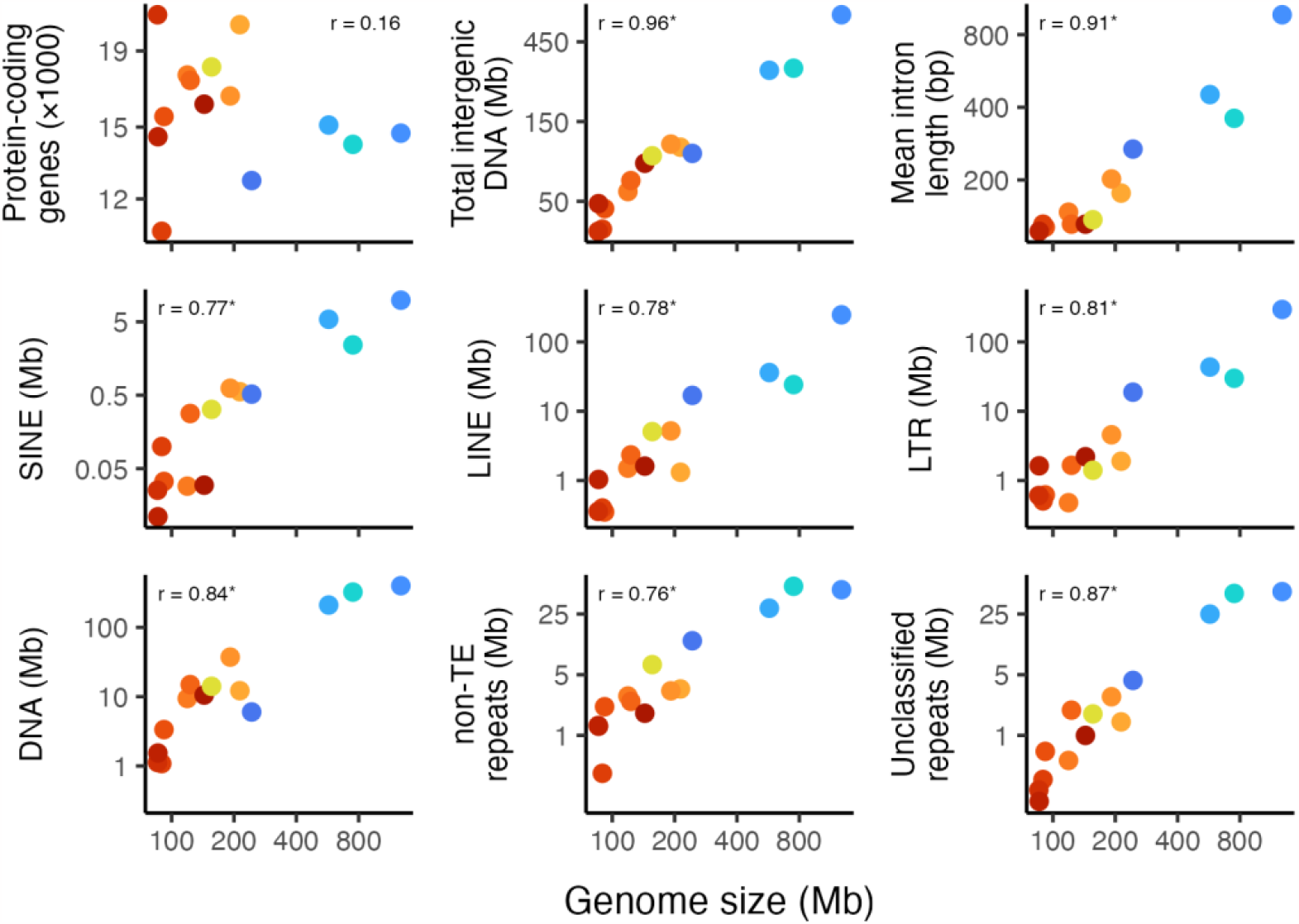
Genome size versus protein-coding genes, intergenic content, mean intron length, and repeat-element content by class for dipterans in our phylogeny. Point colors and data transformations for all panels is as described in fig. 1. Numbers in each panel indicate the estimate of the Pearson correlation coefficient between each variable and genome size using cor_phylo, and ‘*’ indicates the correlation’s 95% confidence interval did not overlap zero (table S2).

### Gene family evolution

Of the 19,860 total phylogenetic Hierarchical Orthologous Groups (HOGs) output from OrthoFinder and used in CAFE for gene family evolution analysis, 9,058 were present at the root of the phylogeny, 736 changed significantly (*P* < 0.05) across the phylogeny, and six had a significant change (*P* < 0.001) at the node separating Chironomidae from their nearest relative, *C. sonorensis* (table S3). GO terms overrepresented in these HOGs spanned a range of biological processes (fig. 3A). Some of these may relate to the ability of chironomids to tolerate stressful or nutrient-limited environments, such as temperature homeostasis, inflammatory response, blood coagulation, and transport of dehydroascorbic acid (the oxidized form of vitamin A). Other overrepresented GO terms may indicate chironomids evolving in response to infectious agents (Toll signaling pathway, response to fungus) and to plant chemical defenses (response to caffeine). Melanization defense response may relate to multiple adaptive pathways since it plays many physiological roles, including desiccation tolerance (Rajpurohit et al. 2008) and immune response (Nakhleh et al. 2017). Similarly, trehalose transport is likely a key stress-associated adaptation because of its diverse functions in many forms of tolerance such as cold and hypoxia (Chen & Haddad 2004; Elbein 2003), as well as protection from desiccation, as in the chironomid *Polypedilum vanderplanki* (Sakurai et al. 2008).

**Figure 3.**
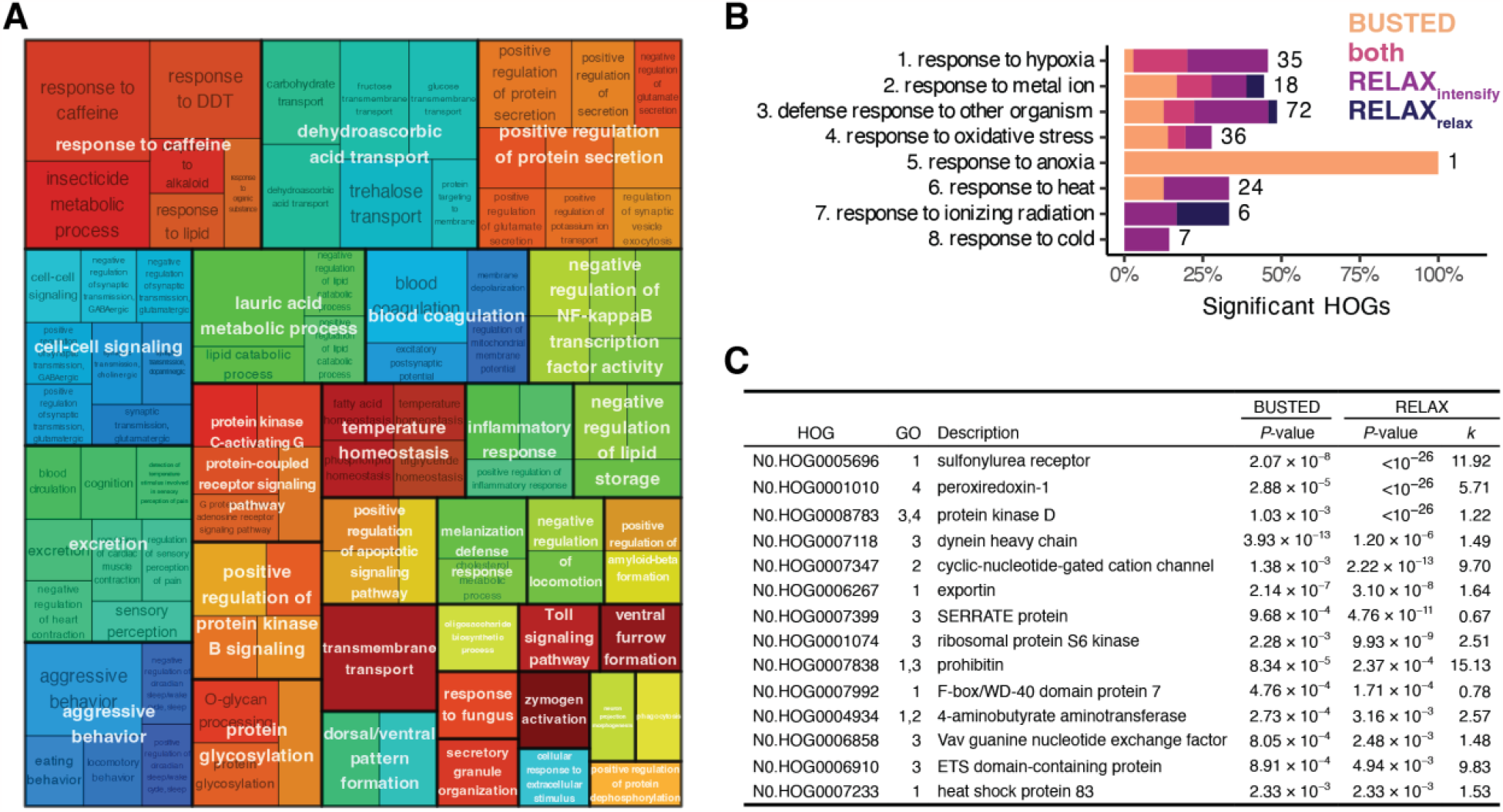
(A) Treemap showing hierarchical structure of GO terms for HOGs that expanded significantly for family Chironomidae. (B) For each targeted GO term related to extreme physiology, percent of associated single-copy HOGs with significant evidence in chironomids for gene-wide positive selection (BUSTED), intensification of selection (RELAX_intensify_), relaxation of selection (RELAX_relax_), or both BUSTED and RELAX (either RELAX_intensify_or RELAX_relax_). Numbers indicate total HOGs per GO term, and some GO terms share HOGs. (C) List of HOGs with evidence for both positive selection and change in selection intensity in chironomids, in order of decreasing evidence for both tests (i.e., *P*_BUSTED_ × *P*_RELAX_). The GO column indicates the GO term(s) listed in panel B associated with each HOG. Parameter *k* indicates the relative selection intensity for Chironomidae compared to outgroups (*k* > 1 means intensified selection, *k* < 1 means relaxed). HOG descriptions were extracted from the representative gene in *Culex quinquefasciatus*.

### Positive selection

We tested for positive selection in 162 single-copy HOGs containing GO terms associated with tolerating stressful environments (fig. 3B; table S4). Using HyPhy’s BUSTED method, 30 (18.5%) HOGs had evidence of gene-wide positive selection in chironomids compared to outgroups. Using HyPhy’s RELAX method, 47 (29.0%) HOGs had evidence for relaxation or intensification of selection in chironomids, with most (41) of these indicating intensification. Fourteen (8.6%) HOGs were significant for both tests, only two of which had evidence for relaxed (instead of intensified) selection. Most of these HOGs were associated with ‘defense response to other organism’ (7) and/or ‘response to hypoxia’ (6) GO terms. The genes with the strongest evidence for positive, intensified selection for Chironomidae were sulfonylurea receptor, peroxiredoxin-1, and protein kinase D. Sulfonylurea receptor is involved in chitin synthesis (Abo-Elghar et al. 2004) and protects against cardiac hypoxic stress in *Drosophila melanogaster* (Akasaka et al. 2006). Peroxiredoxins reduce peroxides and play multiple roles in the inflammation process (Knoops et al. 2016) and can affect lifespan in *D. melanogaster* (Orr et al. 2013). Protein kinase D contributes to oxidative stress signaling and mediating of antioxidant enzyme expression (Storz & Toker 2003; Storz et al. 2005). These genes make especially interesting candidates for further study of the molecular bases of chironomid stress tolerance.

## Materials and Methods

Below is an overview of the methods, and more detailed information can be found in the Supplementary Material online.

### Data sources

In this study, we generated two new reference assemblies, three gene predictions, and four functional annotations. Reference assemblies and genome annotations for the comparative analysis (tables 1 and S5) include previously published data, which were obtained from GenBank, SRA, VectorBase (Amos et al. 2022), or InsectBase (Mei et al. 2022).

### DNA samples, extractions, sequencing

We collected *T. gracilentus* from Lake Mývatn, Iceland (fig. S4A). For long-read ONT sequencing, we extracted high-molecular-weight DNA from a single adult male. ONT library preparation and sequencing were performed by the Roy J. Carver Biotechnology Center, University of Illinois Urbana-Champaign. We also used pooled short-read DNAseq of adults for assembly polishing (as in Steward et al. 2021) and RNAseq of both adults and juveniles to inform gene predictions (fig. S4B). Extractions, library preparation, and sequencing were conducted by the University of Wisconsin–Madison Biotechnology Center.

### Genome assemblies

For our *T. gracilentus* assembly, we first generated multiple assemblies from ONT reads using different assemblers, then combined them into a single best assembly using quickmerge (tables S6, S7). For each step of the assembly, we used BUSCO with the diptera_odb10 dataset to evaluate genome completeness and a custom python script to evaluate contiguity. We estimated the genome size by mapping ONT reads back onto the final assembly using backmap. We generated the assembly for *P. steinenii* using NextDenovo on publicly available ONT sequences (SRA accessions: SRR8180978, SRR3951280, SRR3951285, SRR3951284, SRR3951283). We looked for contamination in both final assemblies using unique 31-mers via sendsketch.sh from bbmap.

### Repeat elements and genome annotations

We described repeat elements for all species in our phylogeny by combining a de novo library of repetitive elements for each using RepeatModeler with a library of dipteran repeats from RepBase. We used RepeatMasker to summarize repeat elements by class and to calculate repeat element divergences. We used BRAKER and the GeneMark-ES Suite to create two sets of gene predictions, one using RNAseq reads and another using the OrthoDB arthropod protein database. We combined them using TSEBRA. We functionally annotated genes using mantis that compares protein sequences to the databases Pfam, KOfam, eggNOG, NCBI’s protein family models, and the Transporter classification database.

### Phylogeny construction and finding orthogroups

To construct the phylogeny, we first extracted amino acid sequences for single-copy orthologs from the diptera_odb10 database using BUSCO. We aligned sequences using MAFFT, concatenated alignments, then used RAxML-NG to generate a maximum likelihood tree and to quantify bootstrapped branch support. We created a time-calibrated tree by combining the ML tree with fossil data from paleobiodb.org (queried on 5 July 2022) and previous time estimates using MCMCTree. We defined five calibration points: (a) 238.5 Ma minimum and 295.4 Ma maximum for the root (Benton et al. 2009), (b) 242.0 Ma minimum for the superfamily Chironomoidea (Lukashevich et al. 2010), (c) 201.3 Ma minimum for family Chironomidae (Krzeminski & Jarzembowski 1999), (d) 93.5 Ma minimum for subfamily Chironominae (Giłka et al. 2022), and (e) 33.9 Ma minimum for the portion of subfamily Orthocladiinae containing genera *Belgica* and *Clunio* (Zelentsov et al. 2012). These estimates informed the parameters for the divergence time sampling distributions in MCMCTree (see table S8 for specific details). MCMCTree was implemented using CODEML to inform the overall substitution rate and the “data-driven birth–death” method (Tao et al. 2021) to inform priors for the speciation birth–death process.

We used OrthoFinder to identify phylogenetic HOGs. We input each species’ protein sets (filtering for longest isoform per gene) and the time-calibrated species tree into OrthoFinder and used the set of HOGs for the root of the phylogeny for downstream analyses. We removed *Clunio marinus* from any analyses using HOGs because only 58.5% of its genes were assigned to orthogroups.

### Features associated with genome size

We looked for associations between genome size and four genomic features in chironomids: protein-coding genes, intergenic sequences, introns, and repeat elements. We first tested for significant differences in each feature (including genome size) between chironomids and other dipterans to assess whether any associations likely pertain to chironomids specifically. For these tests, we grouped *C. sonorensis* (family Ceratopogonidae) with chironomids because exploratory plots made it clear that it shared similar genomic features with Chironomidae. We used phylogenetic linear regressions via the R package phylolm (Ho & Ane 2014). We next tested for whether each feature was correlated with genome size to ascertain whether any changes that occurred likely contributed to genome compaction using the cor_phylo function in the phyr package (Li et al. 2020) that accounts for phylogenetic covariance.

### Gene family evolution

We used CAFE to identify HOGs that significantly expanded (*P* < 0.001) at the node separating Chironomidae from their most recent common ancestor. We used the enricher function in clusterProfiler to find enriched GO terms in our set of HOGs. We then used rrvgo to reduce the redundancy of the set of enriched GOs and to summarize them using a treemap.

### Positive selection

We used HyPhy to test for positive selection in Chironomidae in single-copy HOGs that were associated with any of our *a priori* list of GO terms associated with tolerance to extreme habitats (table S4). We labelled Chironomidae on our tree to define our foreground branches, then for each HOG, we tested (1) whether positive selection occurred for any chironomids using HyPhy’s BUSTED method (Murrell et al. 2015) and (2) whether selection intensified or relaxed for chironomids using HyPhy’s RELAX method (Wertheim et al. 2015). We corrected *P*-values for multiple comparisons using the Benjamini–Yekutieli procedure.

## Data availability

All reference assemblies and raw sequencing data are archived on NCBI under BioProject number PRJNA1044157. Data resulting from analyses are on Zenodo at https://dx.doi.org/10.5281/zenodo.10366330, and all scripts to run the analyses are archived on Zenodo at https://dx.doi.org/10.5281/zenodo.10367770.

## Acknowledgements

This work was supported in part by the American Society of Naturalists Student Research Award (L.A.N.), the National Science Foundation grant DEB 1556208 (A.R.I), and the Steenbock Professorship in Biological Sciences (A.R.I.). We thank the Roy J. Carver Biotechnology Center, University of Illinois Urbana-Champaign for providing guidance on ONT sequencing, and the DNA Sequencing Facility and Gene Expression Center at the University of Wisconsin–Madison Biotechnology Center for providing consultation and services for the Illumina sequencing. This research was performed using the compute resources and assistance of the UW–Madison Center For High Throughput Computing (CHTC) in the Department of Computer Sciences.

## Literature Cited

Abo-Elghar GE, Fujiyoshi P, Matsumura F. 2004. Significance of the sulfonylurea receptor (SUR) as the target of diflubenzuron in chitin synthesis inhibition in Drosophila melanogaster and Blattella germanica. Insect Biochemistry and Molecular Biology. 34:743–752. doi: 10.1016/j.ibmb.2004.03.009.

Akasaka T et al. 2006. The ATP-sensitive potassium (K _ATP_) channel-encoded dSUR gene is required for Drosophila heart function and is regulated by tinman. Proc. Natl. Acad. Sci. U.S.A. 103:11999–12004. doi: 10.1073/pnas.0603098103.

Amos B et al. 2022. VEuPathDB: the eukaryotic pathogen, vector and host bioinformatics resource center. Nucleic Acids Research. 50:D898–D911. doi: 10.1093/nar/gkab929.

Armitage PD, Cranston PS, Pinder LCV, eds. 1995. The Chironomidae. The Biology and Ecology of Non-biting Midges. Chapman & Hall: London, UK.

Benton MJ, Donoghue PCJ, Asher RJ. 2009. Calibrating and constraining molecular clocks. In: The Timetree of Life. Hedges, SB & Kumar, S, editors. Oxford University Press: New York, NY, USA pp. 35–86.

Bertone MA, Courtney GW, Wiegmann BM. 2008. Phylogenetics and temporal diversification of the earliest true flies (Insecta: Diptera) based on multiple nuclear genes. Systematic Entomology. 33:668–687. doi: 10.1111/j.1365-3113.2008.00437.x.

Burmester T, Hankeln T. 2007. The respiratory proteins of insects. Journal of Insect Physiology. 53:285–294. doi: 10.1016/j.jinsphys.2006.12.006.

Chen Q, Haddad GG. 2004. Role of trehalose phosphate synthase and trehalose during hypoxia: from flies to mammals. Journal of Experimental Biology. 207:3125–3129. doi: 10.1242/jeb.01133.

Cong Y, Ye X, Mei Y, He K, Li F. 2022. Transposons and non-coding regions drive the intrafamily differences of genome size in insects. iScience. 25:104873. doi: 10.1016/j.isci.2022.104873.

Cornette R et al. 2015. Chironomid midges (Diptera, Chironomidae) show extremely small genome sizes. Zoological Science. 32:248–254.

Cranston PS, Hardy NB, Morse GE. 2012. A dated molecular phylogeny for the Chironomidae (Diptera). Systematic Entomology. 37:172–188.

Cranston PS, Hardy NB, Morse GE, Puslednik L, McCLUEN SR. 2010. When molecules and morphology concur: the ‘Gondwanan’ midges (Diptera: Chironomidae). Systematic Entomology. 35:636–648. doi: 10.1111/j.1365-3113.2010.00531.x.

Elbein AD. 2003. New insights on trehalose: a multifunctional molecule. Glycobiology. 13:17R –27. doi: 10.1093/glycob/cwg047.

Giłka W et al. 2022. Wanted, tracked down and identified: Mesozoic non-biting midges of the subfamily Chironominae (Chironomidae, Diptera). Zoological Journal of the Linnean Society. 194:874–892. doi: 10.1093/zoolinnean/zlab020.

Gusev O et al. 2010. Anhydrobiosis-associated nuclear DNA damage and repair in the sleeping chironomid: linkage with radioresistance Zhou, Z, editor. PLoS ONE. 5:e14008. doi: 10.1371/journal.pone.0014008.

Gusev O et al. 2014. Comparative genome sequencing reveals genomic signature of extreme desiccation tolerance in the anhydrobiotic midge. Nature Communications. 5:4784.

Ho LST, Ane C. 2014. A linear-time algorithm for Gaussian and non-Gaussian trait evolution models. Systematic Biology. 63:397–408.

Kelley JL et al. 2014. Compact genome of the Antarctic midge is likely an adaptation to an extreme environment. Nature Communications. 5:1–8.

Knoops B, Argyropoulou V, Becker S, Ferté L, Kuznetsova O. 2016. Multiple roles of peroxiredoxins in inflammation. Molecules and Cells. 39:60–64. doi: 10.14348/molcells.2016.2341.

Kohshima S. 1984. A novel cold-tolerant insect found in a Himalayan glacier. Nature. 310:225–227. doi: 10.1038/310225a0.

Krzeminski W, Jarzembowski E. 1999. Aenne triassica sp. n., the oldest representative of the family Chironomidae (Insecta: Diptera). Polskie Pismo Entomologiczne. 68:445–449.

Lee RE et al. 2006. Rapid cold-hardening increases the freezing tolerance of the Antarctic midge Belgica antarctica. Journal of Experimental Biology. 209:399–406. doi: 10.1242/jeb.02001.

Li D, Dinnage R, Nell LA, Helmus MR, Ives AR. 2020. phyr: an R package for phylogenetic species-distribution modelling in ecological communities Price, S, editor. Methods in Ecology and Evolution. 11:1455–1463. doi: 10.1111/2041-210X.13471.

Linevich A. 1963. K biologii komarov semeistva Tendipedidae ‘Biologiya bespozvonochnykh Baikala’. Trudy Limnologicheskogo Instituta. 1:3–48.

Lopez-Martinez G, Elnitsky MA, Benoit JB, Lee RE, Denlinger DL. 2008. High resistance to oxidative damage in the Antarctic midge Belgica antarctica, and developmentally linked expression of genes encoding superoxide dismutase, catalase and heat shock proteins. Insect Biochemistry and Molecular Biology. 38:796–804. doi: 10.1016/j.ibmb.2008.05.006.

Lukashevich ED, Przhiboro AA, Marchal-Papier F, Grauvogel-Stamm L. 2010. The oldest occurrence of immature Diptera (Insecta), Middle Triassic, France. Annales de la Société entomologique de France (N.S.). 46:4–22. doi: 10.1080/00379271.2010.10697636.

Mei Y et al. 2022. InsectBase 2.0: a comprehensive gene resource for insects. Nucleic Acids Research. 50:D1040–D1045. doi: 10.1093/nar/gkab1090.

Murrell B et al. 2015. Gene-wide identification of episodic selection. Molecular Biology and Evolution. 32:1365–1371. doi: 10.1093/molbev/msv035.

Nakhleh J, El Moussawi L, Osta MA. 2017. The melanization response in insect immunity. In: Advances in Insect Physiology.Vol. 52 Elsevier pp. 83–109. doi: 10.1016/bs.aiip.2016.11.002.

Oliver D, Corbet P. 1966. Aquatic habitats in a high arctic locality: the hazen camp study area, Ellesmere Island, NWT. Defense Research Board of Canada.

Orr WC, Radyuk SN, Sohal RS. 2013. Involvement of redox state in the aging of Drosophila melanogaster. Antioxidants & Redox Signaling. 19:788–803. doi: 10.1089/ars.2012.5002.

Rainford JL, Hofreiter M, Nicholson DB, Mayhew PJ. 2014. Phylogenetic distribution of extant richness suggests metamorphosis is a key innovation driving diversification in insects Janke, A, editor. PLoS ONE. 9:e109085. doi: 10.1371/journal.pone.0109085.

Rajpurohit S, Parkash R, Ramniwas S. 2008. Body melanization and its adaptive role in thermoregulation and tolerance against desiccating conditions in drosophilids. Entomological Research. 38:49–60. doi: 10.1111/j.1748-5967.2008.00129.x.

Rinehart JP et al. 2006. Continuous up-regulation of heat shock proteins in larvae, but not adults, of a polar insect. Proc. Natl. Acad. Sci. U.S.A. 103:14223–14227. doi: 10.1073/pnas.0606840103.

Sakurai M et al. 2008. Vitrification is essential for anhydrobiosis in an African chironomid, Polypedilum vanderplanki. Proc. Natl. Acad. Sci. U.S.A. 105:5093–5098. doi: 10.1073/pnas.0706197105.

Shin SC et al. 2019. Nanopore sequencing reads improve assembly and gene annotation of the Parochlus steinenii genome. Scientific reports. 9:1–10.

Steward RA, Okamura Y, Boggs CL, Vogel H, Wheat CW. 2021. The genome of the margined white butterfly (Pieris macdunnoughii): sex chromosome insights and the power of polishing with PoolSeq data Lavrov, D, editor. Genome Biology and Evolution. 13:evab053. doi: 10.1093/gbe/evab053.

Storz P, Döppler H, Toker A. 2005. Protein kinase D mediates mitochondrion-to-nucleus signaling and detoxification from mitochondrial reactive oxygen species. Molecular and Cellular Biology. 25:8520–8530. doi: 10.1128/MCB.25.19.8520-8530.2005.

Storz P, Toker A. 2003. Protein kinase D mediates a stress-induced NF-κB activation and survival pathway. The EMBO Journal. 22:109–120. doi: 10.1093/emboj/cdg009.

Sun X et al. 2021. A chromosome level genome assembly of Propsilocerus akamusi to understand its response to heavy metal exposure. Mol Ecol Resour. 21:1996–2012. doi: 10.1111/1755-0998.13377.

Tao Q, Barba-Montoya J, Kumar S. 2021. Data-driven speciation tree prior for better species divergence times in calibration-poor molecular phylogenies. Bioinformatics. 37:i102–i110. doi: 10.1093/bioinformatics/btab307.

Usher MB, Edwards M. 1984. A dipteran from south of the Antarctic Circle: Belgica antarctica (Chironomidae) with a description of its larva. Biological Journal of the Linnean Society. 23:19–31. doi: 10.1111/j.1095-8312.1984.tb00803.x.

Watanabe M, Kikawada T, Minagawa N, Yukuhiro F, Okuda T. 2002. Mechanism allowing an insect to survive complete dehydration and extreme temperatures. Journal of Experimental Biology. 205:2799–2802. doi: 10.1242/jeb.205.18.2799.

Wertheim JO, Murrell B, Smith MD, Kosakovsky Pond SL, Scheffler K. 2015. RELAX: detecting relaxed selection in a phylogenetic framework. Molecular Biology and Evolution. 32:820–832. doi: 10.1093/molbev/msu400.

Yoshida Y et al. 2022. High quality genome assembly of the anhydrobiotic midge provides insights on a single chromosome-based emergence of extreme desiccation tolerance. NAR Genomics and Bioinformatics. 4:lqac029. doi: 10.1093/nargab/lqac029.

Zelentsov NI, Baranov VA, Perkovsky EE, Shobanov NA. 2012. First records on non-biting midges (Diptera: Chironomidae) from the Rovno amber. Russian Entomological Journal. 21:79–87. doi: 10.15298/rusentj.21.1.10.

Zheng X, Long W, Guo Y, Ma E. 2011. Effects of cadmium exposure on lipid peroxidation and the antioxidant system in fourth-instar larvae of Propsilocerus akamusi (Diptera: Chironomidae) under laboratory conditions. Jnl. Econ. Entom. 104:827–832. doi: 10.1603/EC10109.

Zheng X, Xu Z, Qin G, Wu H, Wei H. 2017. Cadmium exposure on tissue-specific cadmium accumulation and alteration of hemoglobin expression in the 4th-instar larvae of Propsilocerus akamusi (Tokunaga) under laboratory conditions. Ecotoxicology and Environmental Safety. 144:187–192. doi: 10.1016/j.ecoenv.2017.06.019.

