## Supplementary Materials for "Shared features underlying compact genomes and extreme habitat use in chironomid midges"

#### Supplementary Materials and Methods

The full dataset we analyzed included 14 total species from order Diptera, nine of which were from family Chironomidae (table 1). Outside Chironomidae but within infraorder Culicomorpha, we included one species from family Ceratopogonidae (*Culicoides sonorensis*) and three from family Culicidae: *Anopheles stephensi*, *Aedes aegypti*, and *Culex quinquefasciatus*. To more fully represent order Diptera, we also included *Musca domestica*, which is part of suborder Brachycera, family Muscidae. We gathered or created reference assemblies, gene predictions, functional annotations, and repeat libraries for all 14 species. Of these species, we generated new (1) sequencing reads for one (*Tanytarsus gracilentus*), (2) reference assemblies for two (*Parochlus steinenii* and *T. gracilentus*), (3) gene predictions for three (*C. sonorensis*, *P. steinenii*, and *T. gracilentus*), (4) functional annotations for four (*Chironomus riparius*, *C. sonorensis*, *P. steinenii*, and *T. gracilentus*), and (5) repeat libraries for all species. The data we did not generate ourselves we collected from GenBank, Sequence Read

Archive (SRA), VectorBase (Amos et al. 2022; Giraldo-Calderón et al. 2015), or InsectBase (Mei et al. 2022). In the next sections, we describe our data collection in order of use, from sample collection to functional annotations. We then describe how we used these data to construct a time-calibrated phylogeny and analyze genomic features associated with compact genomes and extreme physiology in Chironomidae.

##### ***Tanytarsus gracilentus*** **samples, extractions, and sequencing**

We collected all *Tanytarsus gracilentus* from sites at Lake Mývatn, Iceland (65.5852º, −16.9816º) (fig. S4). DNA samples were stored in 95% ethanol and RNA samples in RNAlater, and all were kept at −20ºC in Iceland. During and after transport to the United States, DNA samples were stored at −80ºC, while RNA samples remained at −20ºC until extraction.

For long-read Oxford Nanopore Technologies (ONT) sequencing, we extracted high-molecular-weight DNA from a single adult male using a previously published protocol (Quick 2018). ONT library preparation and sequencing were performed by the Roy J. Carver Biotechnology Center, University of Illinois Urbana-Champaign. They used 50ng size-selected DNA to create a PCR library with the EXP-PBC001 and LSK-109 kits (Oxford Nanopore Technologies, Oxford, UK) and sequenced the library on one R9.4.1 FLO-MIN106 RevD SpotON flowcell for 72 hours on a MinION II sequencer.

We also used pooled short-read DNAseq for assembly polishing (Steward et al. 2021) and RNAseq to inform gene predictions. Extractions, library preparation, and sequencing were conducted by the University of Wisconsin–Madison Biotechnology Center, with sequencing on an Illumina NovaSeq 6000 (2 × 150bp reads on an S4 flow cell). DNA was extracted from a pool of 7 adult males using a QIAcube HT from QIAGEN (Hilden, Germany), and DNA concentration was verified using the Qubit dsDNA HS Assay Kit (Life Technologies, Grand Island, NY). The sample was then prepared according to the Celero PCR Workflow with Enzymatic Fragmentation (Tecan Genomics, Redwood City, CA). Quality and quantity of the finished library was assessed using an Agilent TapeStation (Agilent, Santa Clara, CA) and Qubit dsDNA HS Assay Kit, respectively. DNAseq libraries were sequenced at ≥ 100× coverage. Two RNAseq libraries were also generated, one from a pool of 10 adult males and another from a pool of 155 first-instar juveniles. Samples were homogenized using a QIAGEN TissueLyser, then RNA was extracted using organic extractions followed by on-column DNase treatment with QIAGEN RNeasy Mini Kit. RNA quality and quantity were assessed using a NanoDrop (Thermo Fisher Scientific, Waltham, MA) and Agilent BioAnalyzer. Libraries were created using the Illumina TruSeq Stranded mRNA library preparation kit and were sequenced to obtain ≥ 100 million reads per sample.

##### ***Tanytarsus gracilentus* genome assembly**

We performed ONT basecalling and adapter trimming using Guppy v5.0.11 and the dna_r9.4.1_450bps_hac.cfg model. For both DNAseq and RNAseq Illumina reads, we used fastp v0.23.2 (Chen et al. 2018) for adapter trimming, polyG tail trimming, and correction in overlapping regions of paired reads. For RNAseq reads, we also used fastp to remove polyA tails, and we followed previous recommendations for mRNA data (MacManes 2014) by applying modest quality trimming (Q ≥ 5) and a read length filter (≥ 50 bp). For mapping reads to assemblies throughout, we used minimap2 v2.24 (Li 2018) for ONT reads, BWA-MEM v0.7.17 (Li 2013) for Illumina DNAseq, and HISAT2 v2.2.1 (Kim et al. 2019) Illumina RNAseq.

Our assembly strategy for *T. gracilentus* consisted of using multiple assemblers, then combining them into a single best assembly using quickmerge v0.3 (Chakraborty et al. 2016) (fig. S5, table S6). For each step of the assembly, we used BUSCO v5.4.3 (Manni et al. 2021) with the diptera_odb10 dataset (Kriventseva et al. 2019) to evaluate genome completeness and a custom python script to evaluate contiguity. To generate initial assemblies from ONT reads, we used NECAT v0.0.1 (Chen et al. 2021), NextDenovo v2.5.0 (Hu et al. 2023), SMARTdenovo v1.0.0 (Liu et al. 2021), and Flye v2.9 (Kolmogorov et al. 2019). Each was followed by ONT polishing: Racon v1.5.0 (3 rounds) (Vaser et al. 2017) then medaka v1.6.0 (https://github.com/nanoporetech/medaka). We assessed assemblies for evidence of heterozygosity-induced duplicate regions via the hist_plot.py script in purge_dups v1.2.5 (Guan et al. 2020), and then used purge_dups to remove repeated regions. We next merged every pairwise permutation of these assemblies using quickmerge. The four best permutations were used in a second round of merges that produced a single best assembly (table S7). We polished this assembly again with ONT reads using medaka, then polished it using pooled Illumina DNAseq reads (with uncalled bases removed using fastp) using NextPolish v1.4.0 (Hu et al. 2020). In addition to evaluating the final assembly completeness using BUSCO, we estimated the genome size by mapping ONT reads back onto the final assembly using backmap v0.5 (Pfenninger et al. 2022) and looked for contamination in the final assembly using sendsketch.sh (default settings) from bbmap v38.96 (Bushnell 2014).

***Parochlus steinenii* reads and genome assembly**

We generated a new reference assembly for *Parochlus steinenii* based on previously published sequencing data (Shin et al. 2019). This species is especially useful because of its place within the subfamily Podonominae, which represents an outgroup with respect to *T. gracilentus* and the other chironomid species with reference assemblies. We used ONT reads (SRA accession: SRR8180978) and paired-end Illumina DNAseq reads (SRR3951280) for genome assembly, and paired-end Illumina RNAseq reads (SRR3951285, SRR3951284, and SRR3951283) for the annotation. Trimming of Illumina reads followed the methods described for *T. gracilentus*. We generated an initial assembly based on ONT reads using NextDenovo v2.5.0. We polished the assembly using 6 rounds of Racon with ONT reads, then using 6 rounds of NextPolish with Illumina DNAseq reads.

##### *Culicoides sonorensis* RNAseq reads

As a member of family Ceratopogonidae, the most closely related family to Chironomidae, *Culicoides sonorensis* is especially informative for understanding the evolutionary history of Chironomidae. To generate gene predictions for *C. sonorensis*, we used paired-end Illumina RNAseq reads from a project designed to boost gene expression under many different environmental conditions (Morales-Hojas et al. 2018). We accessed these reads from the SRA using the following accession codes: ERR637904, ERR637905, ERR637906, ERR637907, ERR637908, ERR637909, ERR637910, ERR637911, ERR637912, ERR637913, and ERR637914.

##### Repeat elements

We described repeat elements for all 14 species' assemblies by first creating a de novo library of repetitive elements for each using RepeatModeler v2.0.3 (Flynn et al. 2020) with the LTR structural finder enabled, then identifying many of the unclassified elements using TEClass v2.1.3 (Abrusán et al. 2009). Next, we combined this custom library with dipteran repeats from the RepBase RepeatMasker edition library v20181026 (Bao et al. 2015). We then soft-masked the reference assembly based on this combined library using RepeatMasker v4.1.2 (Smit et al. 2013). In the species for which we created new gene predictions, we used this soft-masked assembly for all analyses downstream, but for the others, we continued to use the original assemblies that were already soft masked. For all species, we used the annotation (*.out) file from RepeatMasker to summarize repeat elements by class. Next, we calculated repeat element divergences (CpG-adjusted Kimura substitution level) using RepeatMasker's calcDivergenceFromAlign.pl script that takes a RepeatMasker align file as input. We manually parsed the resulting divsum files to create repeat landscape plots.

**Gene predictions and functional annotations**

For the *T. gracilentus*, *P. steinenii*, and *C. sonorensis* genome annotations, we used BRAKER v2.1.6 (Brůna et al. 2021) and the GeneMark-ES Suite v4.71 to create two sets of gene predictions. The first set was based on RNAseq reads using GeneMark-ET (Lomsadze et al. 2014) and AUGUSTUS v3.4.0 (Stanke et al. 2008). The second set was based on the arthropod protein database from OrthoDB v10 using ProtHint v2.6.0, GeneMark-EP (Brůna et al. 2020), and AUGUSTUS. These two sets of predictions were combined to create a single consensus gene set using TSEBRA v1.0.3 (Gabriel et al. 2021) with a preference for genes predicted by RNAseq by using the provided pref_braker1.cfg configuration file. We ran BUSCO on the CDS of the annotations to assess performance of the gene predictions. We functionally annotated genes for *Chironomus riparius*, as well as *T. gracilentus*, *P. steinenii*, and *C. sonorensis*, using mantis v1.5.5 (Queirós et al. 2021). This program uses DIAMOND v2.0.15 (Buchfink et al. 2021) and HMMER v3.3.2 (hmmer.org) to compare protein sequences to five different databases: Pfam (El-Gebali et al. 2019), KOfam (Aramaki et al. 2020), eggNOG (Huerta-Cepas et al. 2019), NCBI's protein family models (NPFM) (Lu et al. 2020), and the Transporter classification database (TCDB) (Saier et al. 2021). Databases were accessed on 12 July 2022.

**Other data sources**

We used reference assemblies from GenBank for seven chironomids (table 1). We also included assemblies from outgroups in families Ceratopogonidae (*Culicoides sonorensis*), Culicidae (*Anopheles stephensi*, *Aedes aegypti*, and *Culex quinquefasciatus*), and Muscidae (*Musca domestica*) (table S5). To find gene predictions and functional annotations for all species except those described in previous sections, we used GenBank, VectorBase (Amos et al. 2022; Giraldo-Calderón et al. 2015), and InsectBase (Mei et al. 2022) (table S5).

**Phylogeny construction**

To construct the phylogenomic tree, we first used BUSCO v5.4.3 to extract the amino acid sequences for all diptera_odb10 proteins present and with only a single copy in each assembly. We removed non-homologous characters from unaligned amino acid sequences separately for each protein present in all assemblies using PREQUAL v1.02 (Whelan et al. 2018), then aligned sequences using MAFFT v7.515 (Katoh & Standley 2013). After concatenating alignments, RAxML-NG v1.1.0 (Kozlov et al. 2019) was used to generate a maximum likelihood (ML) tree (using model LG+I+G4m and with *Musca domestica* as outgroup) and assess branch support via 1,000 bootstrap (Felsenstein 1985) replicates. We created a time-calibrated, ultrametric tree by combining the ML tree with fossil records (queried from paleobiodb.org on 5 July 2022) using the Bayesian MCMCTree program in the PAML v4.10.6 (Yang 2007) package. Before running MCMCTree, we used (1) PAML's CODEML program to estimate the overall substitution rate and (2) the “data-driven birth–death” (ddBD) method (Tao et al. 2021) to calculate priors for the speciation birth–death process. We ran CODEML on the ML tree using three calibration point estimates, which were simplifications compared to those used in MCMCTree (described below): 232.7 million years ago (Ma) for family Chironomidae (Cranston et al. 2012), 137.573 Ma for subfamily Chironominae (Cranston et al. 2012), and 76 Ma separating genera *Belgica* and *Clunio* (timetree.org). The ddBD method was run on the tree output from RelTime-ML (Mello et al. 2017) in MEGA-CC v11.0.13-1 (Tamura et al. 2021) that took the initial ML tree and concatenated protein sequence alignments as inputs. We used the LG model for both CODEML and RelTime-ML.

In MCMCTree, we used a LG+Γ model, ran the MCMC for 20,000 iterations, and ran the program 4 times to check for convergence. For MCMCTree’s substitution rate priors, we used a shape parameter of 2.0 for a relatively diffuse prior and a rate value that caused the Gamma distribution mean to equal the substitution rate estimated by CODEML ($9.0526 \times 10^{-10}$ substitutions per year). We used both fossil evidence and previous time estimates (Benton et al. 2009; Hedges & Kumar 2009; Cranston et al. 2012, 2010) to inform the parameters defining the divergence time sampling distributions in MCMCTree. We defined five calibration points: (a) 238.5 Ma minimum and 295.4 Ma maximum for the root (Benton et al. 2009), (b) 242.0 Ma minimum for the superfamily Chironomoidea (Lukashevich et al. 2010), (c) 201.3 Ma minimum for family Chironomidae (Krzeminski & Jarzembowski 1999), (d) 93.5 Ma minimum for subfamily Chironominae (Giłka et al. 2022), and (e) 33.9 Ma minimum for the portion of subfamily Orthocladiinae containing genera *Belgica* and *Clunio* (Zelentsov et al. 2012). All constraints were soft, and we adjusted the probability of sampling a time that violates the constraint based on our perceived reliability of the bounds. MCMCTree samples minimum constraints with a truncated Cauchy distribution, and we defined the Cauchy parameters to make the mode near our best estimate and the variance proportional to our confidence in this estimate (see table S8 for details on parameter values).

**Finding orthogroups**

We used OrthoFinder v2.5.4 (Emms & Kelly 2019) to identify phylogenetic Hierarchical Orthologous Groups (HOGs) for use in the analyses for intron sizes, gene family evolution, and positive selection. We first filtered all species' protein sequences to only include the longest isoform for each protein-coding gene. We input these filtered protein sets and the time-calibrated species tree from MCMCTree into OrthoFinder and used the set of HOGs for the root of the phylogeny for downstream analyses. We removed *Clunio marinus* from any analyses using HOGs because only 58.5% of its genes were assigned to an orthogroup by OrthoFinder, compared to >90% for all other species.

**Features associated with genome size**

We looked for associations between genome size and four genomic features in chironomids: protein-coding genes, intergenic sequences, introns, and repeat elements. We estimated the number of protein-coding genes and total intergenic sequence length from the assemblies' *.gff files, using the longest isoform for each gene for intergenic sequences. Repeat elements were extracted from all species' RepeatMasker annotation (*.out) files, and we aggregated all non-TE elements (rolling-circles, small RNA, satellites, simple repeats, and low complexity) into one category for analysis. We calculated average intron length based only on genes matching to HOGs that had all species represented and that contained < 4 genes per species (Martín-Durán et al. 2021). We $\log_{10}$-transformed these variables because they were highly right skewed, with intron lengths being transformed before calculating the mean.

We first tested for significant differences in each feature (including genome size) between chironomids and other dipterans to assess whether any associations likely pertain to the specific evolutionary history of chironomids. For these tests, we grouped *Culicoides sonorensis* (family Ceratopogonidae) with chironomids because exploratory plots made it clear that it shared similar genomic traits with its most recent common ancestors in our phylogeny: family Chironomidae. We used phylogenetic linear regressions and tested for significance of a binary variable indicating whether the species belonged to families Ceratopogonidae or Chironomidae. We used the phylolm v2.6.2 (Ho & Ane 2014) package in R v4.3.0 (R Core Team 2023) and ran models using an Ornstein-Uhlenbeck process with a fixed state estimated at the root of the phylogeny. We next tested for whether each feature was correlated with genome size to ascertain whether any changes that occurred likely contributed to genome compaction in Chironomidae. We used the cor_phylo function in the R package phyr v1.1.2 (Li et al. 2020) to compute the Pearson correlation coefficient between features and genome size while accounting for phylogenetic covariance. We constrained the phylogenetic signal to be between 0 and 1 when not doing so resulted in the covariance matrix not being positive definite, which prevented bootstrapping. For both sets of analyses, we used parametric bootstrapping (2,000 repetitions) to compute 95% confidence intervals for the parameters of interest, and we scaled the phylogeny to a maximum depth of 1 for easier interpretation because $\alpha$ scales with phylogeny depth (Ives & Garland 2010).

#### Gene family evolution

We used CAFE v5.0.0 (Mendes et al. 2020) to analyze gene family evolution along our phylogeny, looking specifically for families that expanded for Chironomidae. After CAFE removed families not present at the root of the phylogeny, we analyzed 9,058 total HOGs across our 13 species. We used a Poisson root frequency distribution and a Gamma model with 8 categories; we used 8 because from testing, this is lowest value where the changes in the negative log likelihood began to subside. From the CAFE output, we filtered for HOGs that significantly expanded ($P < 0.001$) at the node separating Chironomidae from their most recent common ancestor. To better understand the functions of these expanded HOGs, we connected Gene Ontology (GO) terms from our functional annotations to each HOG based on all genes from all species that matched to it. We then used the enricher function in the R package clusterProfiler v4.8.2 (Wu et al. 2021) to find enriched GO terms in our set of HOGs, using a Benjamini–Hochberg adjusted *P*-value cutoff of 0.1 and minimum gene size of 1. Next, we used the R package rrvgo v1.12.2 (Sayols 2023)—based on the online tool REVIGO (Supek et al. 2011)—to reduce the redundancy of the set of enriched "biological process" GO terms based on the *Drosophila melanogaster*-derived database org.Dm.eg.db v3.17.0 (Carlson 2023b) and to summarize these GO terms using a treemap.

#### Positive selection

We used HyPhy v2.5.52 (Murrell et al. 2015; Wertheim et al. 2015) to test for positive selection in Chironomidae. We first filtered for HOGs that contained exactly one copy per species and that were associated with at least one GO term (including all offspring) from our *a priori* list of terms associated with tolerance to extreme habitats (table S4). We identified offspring GO terms using GO.db v3.17.0 (Carlson 2023a). Because HyPhy requires codon-aware sequence alignments, we extracted the coding sequence (CDS) for the longest isoform per gene and used HyPhy's codon-msa tool (https://github.com/veg/hyphy-analyses) and MAFFT for the alignments for each HOG. For 12 HOGs, alignments failed in one or more species, so these HOGs were removed from our analyses. Next, we labelled Chironomidae on our tree using phylotree.js (Shank et al. 2018) so that we could use chironomids as the test branches (with the others being background) for our analyses. For each HOG, we tested (1) whether positive selection occurred for any chironomids using HyPhy's BUSTED method (Murrell et al. 2015) and (2) whether selection intensified or relaxed for chironomids compared to outgroups using HyPhy's RELAX method (Wertheim et al. 2015). The resulting set of *P*-values for each test across all HOGs were corrected for multiple comparisons using the Benjamini–Yekutieli procedure (Benjamini & Yekutieli 2001) under arbitrary dependence assumptions and using a false discovery rate of 0.10.

### References

Abrusán G, Grundmann N, DeMester L, Makalowski W. 2009. TEclass—a tool for automated classification of unknown eukaryotic transposable elements. Bioinformatics. 25:1329–1330. doi: 10.1093/bioinformatics/btp084.

Amos B et al. 2022. VEuPathDB: the eukaryotic pathogen, vector and host bioinformatics resource center. Nucleic Acids Research. 50:D898–D911. doi: 10.1093/nar/gkab929.

Aramaki T et al. 2020. KofamKOALA: KEGG Ortholog assignment based on profile HMM and adaptive score threshold Valencia, A, editor. Bioinformatics. 36:2251–2252. doi: 10.1093/bioinformatics/btz859.

Bao W, Kojima KK, Kohany O. 2015. Repbase update, a database of repetitive elements in eukaryotic genomes. Mobile DNA. 6:11. doi: 10.1186/s13100-015-0041-9.

Behura SK et al. 2011. Complete sequences of mitochondria genomes of *Aedes aegypti* and *Culex quinquefasciatus* and comparative analysis of mitochondrial DNA fragments inserted in the nuclear genomes. Insect Biochemistry and Molecular Biology. 41:770–777. doi: 10.1016/j.ibmb.2011.05.006.

Benjamini Y, Yekutieli D. 2001. The control of the false discovery rate in multiple testing under dependency. Ann. Statist. 29. doi: 10.1214/aos/1013699998.

Benton MJ, Donoghue PCJ, Asher RJ. 2009. Calibrating and constraining molecular clocks. In: The Timetree of Life. Hedges, SB & Kumar, S, editors. Oxford University Press: New York, NY, USA pp. 35–86.

Brůna T, Hoff KJ, Lomsadze A, Stanke M, Borodovsky M. 2021. BRAKER2: automatic eukaryotic genome annotation with GeneMark-EP+ and AUGUSTUS supported by a protein database. NAR Genomics and Bioinformatics. 3:lqaa108. doi: 10.1093/nargab/lqaa108.

Brůna T, Lomsadze A, Borodovsky M. 2020. GeneMark-EP+: eukaryotic gene prediction with self-training in the space of genes and proteins. NAR Genomics and Bioinformatics. 2:lqaa026. doi: 10.1093/nargab/lqaa026.

Buchfink B, Reuter K, Drost H-G. 2021. Sensitive protein alignments at tree-of-life scale using DIAMOND. Nat Methods. 18:366–368. doi: 10.1038/s41592-021-01101-x.

Bushnell B. 2014. BBMap: a fast, accurate, splice-aware aligner. https://www.osti.gov/biblio/1241166.

Carlson M. 2023a. GO.db: a set of annotation maps describing the entire Gene Ontology.

Carlson M. 2023b. org.Dm.eg.db: genome wide annotation for fly.

Chakraborty M et al. 2021. Hidden genomic features of an invasive malaria vector, *Anopheles stephensi*, revealed by a chromosome-level genome assembly. BMC Biol. 19:28. doi: 10.1186/s12915-021-00963-z.

Chakraborty M, Baldwin-Brown JG, Long AD, Emerson JJ. 2016. Contiguous and accurate *de novo* assembly of metazoan genomes with modest long read coverage. Nucleic Acids Res. gkw654. doi: 10.1093/nar/gkw654.

Chen S, Zhou Y, Chen Y, Gu J. 2018. fastp: an ultra-fast all-in-one FASTQ preprocessor. Bioinformatics. 34:i884–i890. doi: 10.1093/bioinformatics/bty560.

Chen Y et al. 2021. Efficient assembly of nanopore reads via highly accurate and intact error correction. Nat Commun. 12:60. doi: 10.1038/s41467-020-20236-7.

Cranston PS, Hardy NB, Morse GE. 2012. A dated molecular phylogeny for the Chironomidae (Diptera). Systematic Entomology. 37:172–188.

Cranston PS, Hardy NB, Morse GE, Puslednik L, McCLUEN SR. 2010. When molecules and morphology concur: the 'Gondwanan' midges (Diptera: Chironomidae). Systematic Entomology. 35:636–648. doi: 10.1111/j.1365-3113.2010.00531.x.

El-Gebali S et al. 2019. The Pfam protein families database in 2019. Nucleic Acids Research. 47:D427–D432. doi: 10.1093/nar/gky995.

Emms DM, Kelly S. 2019. OrthoFinder: phylogenetic orthology inference for comparative genomics. Genome Biol. 20:238. doi: 10.1186/s13059-019-1832-y.

Felsenstein J. 1985. Confidence limits on phylogenies: an approach using the bootstrap. Evolution. 39:783–791.

Flynn JM et al. 2020. RepeatModeler2 for automated genomic discovery of transposable element families. Proc. Natl. Acad. Sci. U.S.A. 117:9451–9457. doi: 10.1073/pnas.1921046117.

Gabriel L, Hoff KJ, Brůna T, Borodovsky M, Stanke M. 2021. TSEBRA: transcript selector for BRAKER. BMC Bioinformatics. 22:566. doi: 10.1186/s12859-021-04482-0.

Giłka W et al. 2022. Wanted, tracked down and identified: Mesozoic non-biting midges of the subfamily Chironominae (Chironomidae, Diptera). Zoological Journal of the Linnean Society. 194:874–892. doi: 10.1093/zoolinnean/zlab020.

Giraldo-Calderón GI et al. 2015. VectorBase: an updated bioinformatics resource for invertebrate vectors and other organisms related with human diseases. Nucleic Acids Research. 43:D707–D713. doi: 10.1093/nar/gku1117.

Guan D et al. 2020. Identifying and removing haplotypic duplication in primary genome assemblies Valencia, A, editor. Bioinformatics. 36:2896–2898. doi: 10.1093/bioinformatics/btaa025.

Hedges SB, Kumar S, eds. 2009. *The Timetree of Life*. Oxford University Press: New York, NY, USA.

Ho LST, Ane C. 2014. A linear-time algorithm for Gaussian and non-Gaussian trait evolution models. Systematic Biology. 63:397–408.

Hu J et al. 2023. An efficient error correction and accurate assembly tool for noisy long reads. doi: 10.1101/2023.03.09.531669.

Hu J, Fan J, Sun Z, Liu S. 2020. NextPolish: a fast and efficient genome polishing tool for long-read assembly Berger, B, editor. Bioinformatics. 36:2253–2255. doi: 10.1093/bioinformatics/btz891.

Huerta-Cepas J et al. 2019. eggNOG 5.0: a hierarchical, functionally and phylogenetically annotated orthology resource based on 5090 organisms and 2502 viruses. Nucleic Acids Research. 47:D309–D314. doi: 10.1093/nar/gky1085.

Ives AR, Garland T. 2010. Phylogenetic logistic regression for binary dependent variables. Systematic Biology. 59:9–26. doi: 10.1093/sysbio/syp074.

Kaiser TS et al. 2016. The genomic basis of circadian and circalunar timing adaptations in a midge. Nature. 540:69–73.

Katoh K, Standley DM. 2013. MAFFT Multiple Sequence Alignment Software Version 7: Improvements in Performance and Usability. Molecular Biology and Evolution. 30:772–780. doi: 10.1093/molbev/mst010.

Kelley JL et al. 2014. Compact genome of the Antarctic midge is likely an adaptation to an extreme environment. Nature Communications. 5:1–8.

Kim D, Paggi JM, Park C, Bennett C, Salzberg SL. 2019. Graph-based genome alignment and genotyping with HISAT2 and HISAT-genotype. Nat Biotechnol. 37:907–915. doi: 10.1038/s41587-019-0201-4.

Kolmogorov M, Yuan J, Lin Y, Pevzner PA. 2019. Assembly of long, error-prone reads using repeat graphs. Nat Biotechnol. 37:540–546. doi: 10.1038/s41587-019-0072-8.

Kozlov AM, Darriba D, Flouri T, Morel B, Stamatakis A. 2019. RAxML-NG: a fast, scalable and user-friendly tool for maximum likelihood phylogenetic inference Wren, J, editor. Bioinformatics. 35:4453–4455. doi: 10.1093/bioinformatics/btz305.

Kriventseva EV et al. 2019. OrthoDB v10: sampling the diversity of animal, plant, fungal, protist, bacterial and viral genomes for evolutionary and functional annotations of orthologs. Nucleic Acids Research. 47:D807–D811. doi: 10.1093/nar/gky1053.

Krzeminski W, Jarzembowski E. 1999. *Aenne triassica* sp. n., the oldest representative of the family Chironomidae (Insecta: Diptera). Polskie Pismo Entomologiczne. 68:445–449.

Kutsenko A et al. 2014. The *Chironomus tentans* genome sequence and the organization of the Balbiani ring genes. BMC Genomics. 15:819.

Li D, Dinnage R, Nell LA, Helmus MR, Ives AR. 2020. phyr: an R package for phylogenetic species-distribution modelling in ecological communities Price, S, editor. Methods in Ecology and Evolution. 11:1455–1463. doi: 10.1111/2041-210X.13471.

Li H. 2013. Aligning sequence reads, clone sequences and assembly contigs with BWA-MEM. arXiv. https://arxiv.org/abs/1303.3997.

Li H. 2018. Minimap2: pairwise alignment for nucleotide sequences Birol, I, editor. Bioinformatics. 34:3094–3100. doi: 10.1093/bioinformatics/bty191.

Liu H, Wu S, Li A, Ruan J. 2021. SMARTdenovo: a de novo assembler using long noisy reads. Gigabyte. 2021:1–9. doi: 10.46471/gigabyte.15.

Lomsadze A, Burns PD, Borodovsky M. 2014. Integration of mapped RNA-Seq reads into automatic training of eukaryotic gene finding algorithm. Nucleic Acids Research. 42:e119–e119. doi: 10.1093/nar/gku557.

Lu S et al. 2020. CDD/SPARCLE: the conserved domain database in 2020. Nucleic Acids Research. 48:D265–D268. doi: 10.1093/nar/gkz991.

Lukashevich ED, Przhiboro AA, Marchal-Papier F, Grauvogel-Stamm L. 2010. The oldest occurrence of immature Diptera (Insecta), Middle Triassic, France. Annales de la Société entomologique de France (N.S.). 46:4–22. doi: 10.1080/00379271.2010.10697636.

MacManes MD. 2014. On the optimal trimming of high-throughput mRNA sequence data. Front. Genet. 5. doi: 10.3389/fgene.2014.00013.

Manni M, Berkeley MR, Seppey M, Simão FA, Zdobnov EM. 2021. BUSCO update: novel and streamlined workflows along with broader and deeper phylogenetic coverage for scoring of eukaryotic, prokaryotic, and viral genomes Kelley, J, editor. Molecular Biology and Evolution. 38:4647–4654. doi: 10.1093/molbev/msab199.

Martín-Durán JM et al. 2021. Conservative route to genome compaction in a miniature annelid. Nat Ecol Evol. 5:231–242. doi: 10.1038/s41559-020-01327-6.

Matthews BJ et al. 2018. Improved reference genome of *Aedes aegypti* informs arbovirus vector control. Nature. 563:501–507. doi: 10.1038/s41586-018-0692-z.

Mei Y et al. 2022. InsectBase 2.0: a comprehensive gene resource for insects. Nucleic Acids Research. 50:D1040–D1045. doi: 10.1093/nar/gkab1090.

Mello B, Tao Q, Tamura K, Kumar S. 2017. Fast and accurate estimates of divergence times from big data. Mol Biol Evol. 34:45–50. doi: 10.1093/molbev/msw247.

Mendes FK, Vanderpool D, Fulton B, Hahn MW. 2020. CAFE 5 models variation in evolutionary rates among gene families. Bioinformatics. 36:5516–5518. doi: 10.1093/bioinformatics/btaa1022.

Morales-Hojas R et al. 2018. The genome of the biting midge *Culicoides sonorensis* and gene expression analyses of vector competence for Bluetongue virus. BMC Genomics. 19:624.

Murrell B et al. 2015. Gene-wide identification of episodic selection. Molecular Biology and Evolution. 32:1365–1371. doi: 10.1093/molbev/msv035.

Pfenninger M, Schönnenbeck P, Schell T. 2022. ModEst: Accurate estimation of genome size from next generation sequencing data. Molecular Ecology Resources. 22:1454–1464. doi: 10.1111/1755-0998.13570.

Queirós P, Delogu F, Hickl O, May P, Wilmes P. 2021. Mantis: flexible and consensus-driven genome annotation. GigaScience. 10:giab042. doi: 10.1093/gigascience/giab042.

Quick J. 2018. Ultra-long read sequencing protocol for RAD004 v3. doi: 10.17504/protocols.io.mrxc57n.

R Core Team. 2023. R: A Language and Environment for Statistical Computing. https://www.r-project.org/.

Saier MH et al. 2021. The Transporter Classification Database (TCDB): 2021 update. Nucleic Acids Research. 49:D461–D467. doi: 10.1093/nar/gkaa1004.

Sayols S. 2023. rrvgo: a Bioconductor package for interpreting lists of Gene Ontology terms. microPublication Biology. doi: 10.17912/MICROPUB.BIOLOGY.000811.

Scott JG et al. 2014. Genome of the house fly, *Musca domestica* L., a global vector of diseases with adaptations to a septic environment. Genome Biol. 15:466. doi: 10.1186/s13059-014-0466-3.

Shaikhutdinov NM et al. 2023. Population genomics of two closely related anhydrobiotic midges reveals differences in adaptation to extreme desiccation Pfeifer, S, editor. Genome Biology and Evolution. 15:evad169. doi: 10.1093/gbe/evad169.

Shank SD, Weaver S, Kosakovsky Pond SL. 2018. phylotree.js - a JavaScript library for application development and interactive data visualization in phylogenetics. BMC Bioinformatics. 19:276. doi: 10.1186/s12859-018-2283-2.

Shin SC et al. 2019. Nanopore sequencing reads improve assembly and gene annotation of the *Parochlus steinenii* genome. Scientific reports. 9:1–10.

Smit A, Hubley R, Green P. 2013. *RepeatMasker Open-4.0*. http://www.repeatmasker.org.

Stanke M, Diekhans M, Baertsch R, Haussler D. 2008. Using native and syntenically mapped cDNA alignments to improve de novo gene finding. Bioinformatics. 24:637–644. doi: 10.1093/bioinformatics/btn013.

Steward RA, Okamura Y, Boggs CL, Vogel H, Wheat CW. 2021. The genome of the margined white butterfly (*Pieris macdunnoughii*): sex chromosome insights and the power of polishing with PoolSeq data Lavrov, D, editor. Genome Biology and Evolution. 13:evab053. doi: 10.1093/gbe/evab053.

Sun X et al. 2021. A chromosome level genome assembly of *Propsilocerus akamusi* to understand its response to heavy metal exposure. Mol Ecol Resour. 21:1996–2012. doi: 10.1111/1755-0998.13377.

Supek F, Bošnjak M, Škunca N, Šmuc T. 2011. REVIGO summarizes and visualizes long lists of Gene Ontology terms Gibas, C, editor. PLoS ONE. 6:e21800. doi: 10.1371/journal.pone.0021800.

Tamura K, Stecher G, Kumar S. 2021. MEGA11: molecular evolutionary genetics analysis version 11 Battistuzzi, FU, editor. Molecular Biology and Evolution. 38:3022–3027. doi: 10.1093/molbev/msab120.

Tao Q, Barba-Montoya J, Kumar S. 2021. Data-driven speciation tree prior for better species divergence times in calibration-poor molecular phylogenies. Bioinformatics. 37:i102–i110. doi: 10.1093/bioinformatics/btab307.

Vaser R, Sović I, Nagarajan N, Šikić M. 2017. Fast and accurate de novo genome assembly from long uncorrected reads. Genome Res. 27:737–746. doi: 10.1101/gr.214270.116.

Wertheim JO, Murrell B, Smith MD, Kosakovsky Pond SL, Scheffler K. 2015. RELAX: detecting relaxed selection in a phylogenetic framework. Molecular Biology and Evolution. 32:820–832. doi: 10.1093/molbev/msu400.

Whelan S, Irisarri I, Burki F. 2018. PREQUAL: detecting non-homologous characters in sets of unaligned homologous sequences Hancock, J, editor. Bioinformatics. doi: 10.1093/bioinformatics/bty448.

Wu T et al. 2021. clusterProfiler 4.0: a universal enrichment tool for interpreting omics data. The Innovation. 2:100141. doi: 10.1016/j.xinn.2021.100141.

Yang Z. 2007. PAML 4: phylogenetic analysis by maximum likelihood. Molecular Biology and Evolution. 24:1586–1591. doi: 10.1093/molbev/msm088.

Yoshida Y et al. 2022. High quality genome assembly of the anhydrobiotic midge provides insights on a single chromosome-based emergence of extreme desiccation tolerance. NAR Genomics and Bioinformatics. 4:lqac029. doi: 10.1093/nargab/lqac029.

Zelentsov NI, Baranov VA, Perkovsky EE, Shobanov NA. 2012. First records on non-biting midges (Diptera: Chironomidae) from the Rovno amber. Russian Entomological Journal. 21:79–87. doi: 10.15298/rusentj.21.1.10.

### Supplementary Figures and Tables

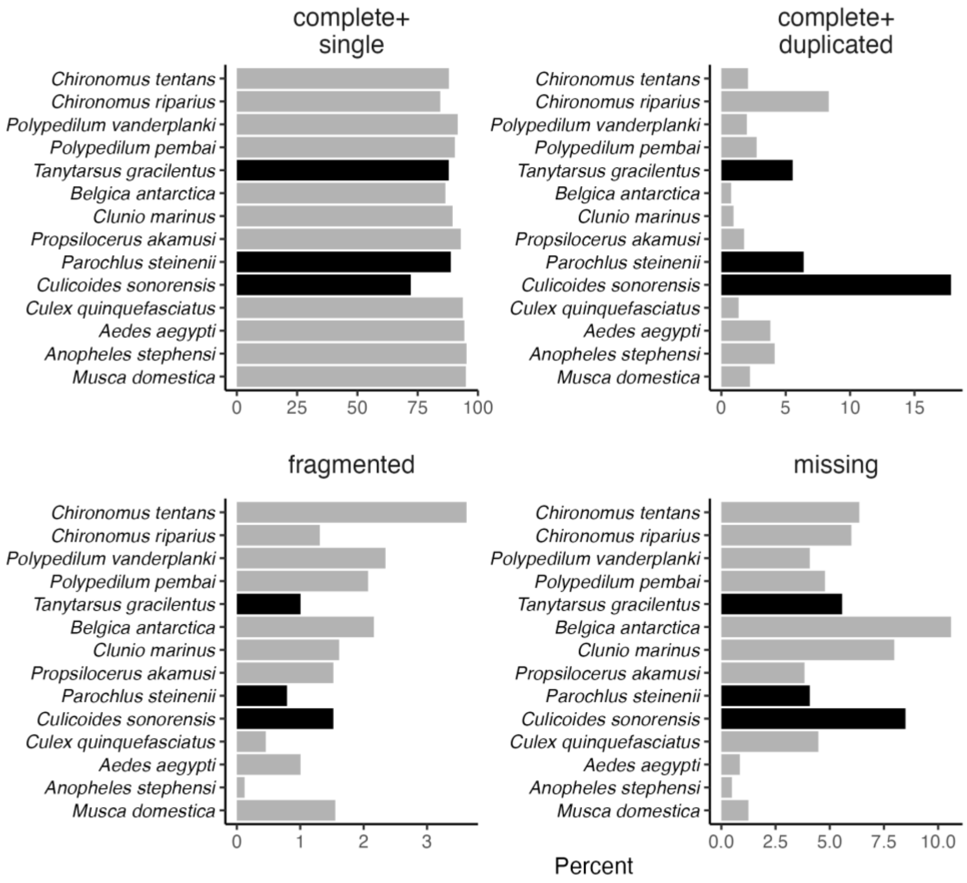

**Figure S1.** Transcriptome BUSCO completeness for all species our phylogeny. Species are ordered as they appear in the phylogeny (figs. 1A, S2), and the three species whose gene predictions were created here are shown in black.

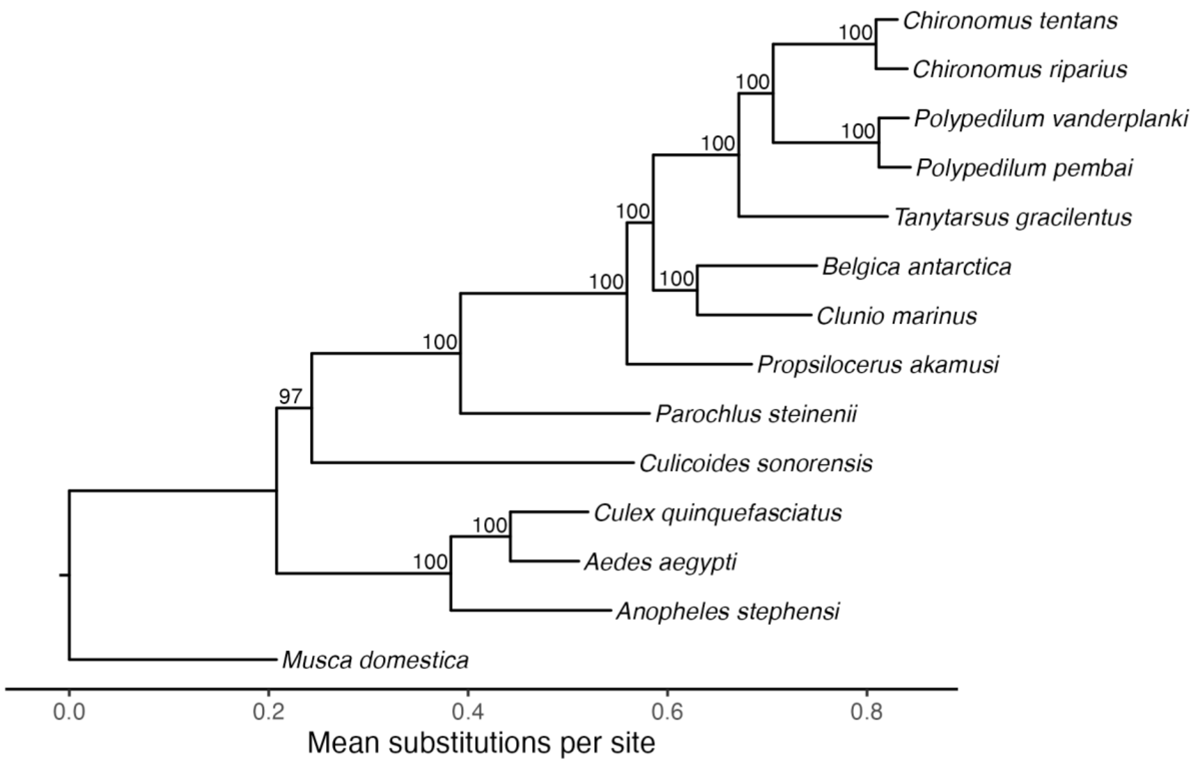

**Figure S2.** Maximum likelihood phylogenomic tree showing relationships among chironomids and outgroups within the order Diptera. Numbers near nodes indicate percent bootstrap support.

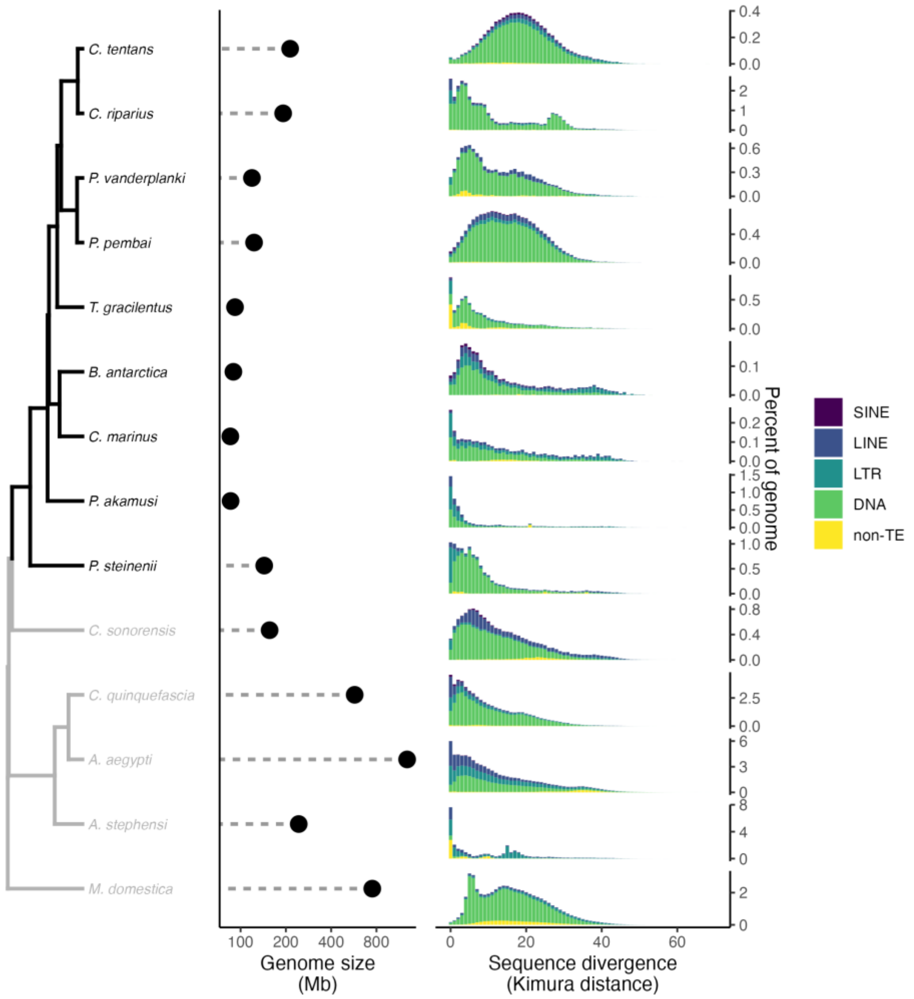

**Figure S3.** Distributions of repeat element divergences calculated using the CpG-adjusted Kimura substitution model (right panel) for all species in our phylogeny, where color indicates the repeat class. The phylogeny (left panel) has chironomids in black and others in gray.

**A**

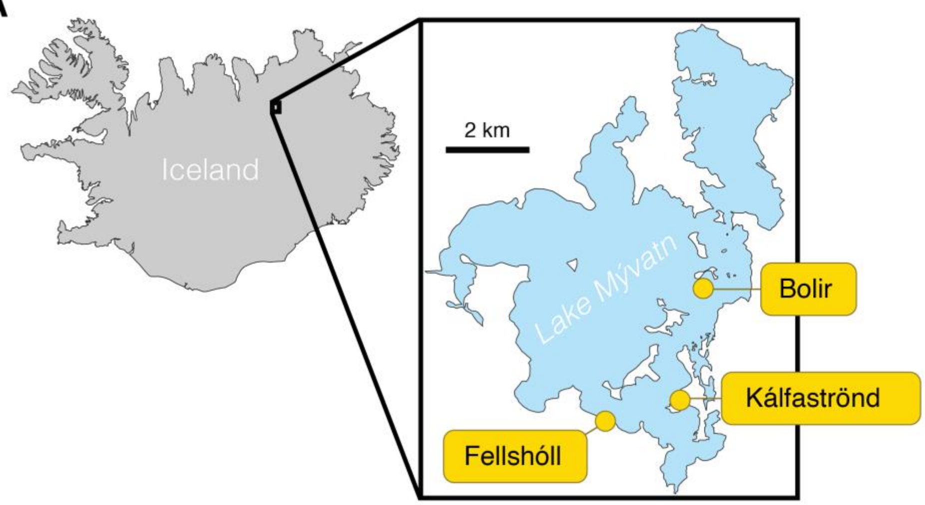

**B**

| Individuals | Life stage | Date collected | Location | Reads type | Platform* | Use(s) |
|---|---|---|---|---|---|---|
| 1 | adult | 10 Jun 2018 | Fellshóll | long DNA | ONT | assembly & polishing |
| 7 | adult | 21 Jun 2019 | Kálfaströnd | 2×150bp DNA | Illumina | polishing |
| 10 | adult | 9 Aug 2021 | Kálfaströnd | 2×150bp RNA | Illumina | annotation |
| 155 | juvenile | 12–13 Aug 2018 | Bolir | 2×150bp RNA | Illumina | annotation |

* "ONT" = Oxford Nanopore MinION II and "Illumina" = Illumina NovaSeq 6000

**Figure S4.** Map and descriptions of sequenced samples. (A) Sampled *Tanytarsus gracilentus* were collected from one of three locations at Lake Mývatn, Iceland. (B) Descriptions of the four sequencing libraries used in this study, including number of *T. gracilentus* individuals in the pool, life stage of the individuals, date collected, location of collection (names coincide with part A), type of reads (long vs 2×150bp, DNA vs RNA), sequencing platform, and step(s) for which they were used.

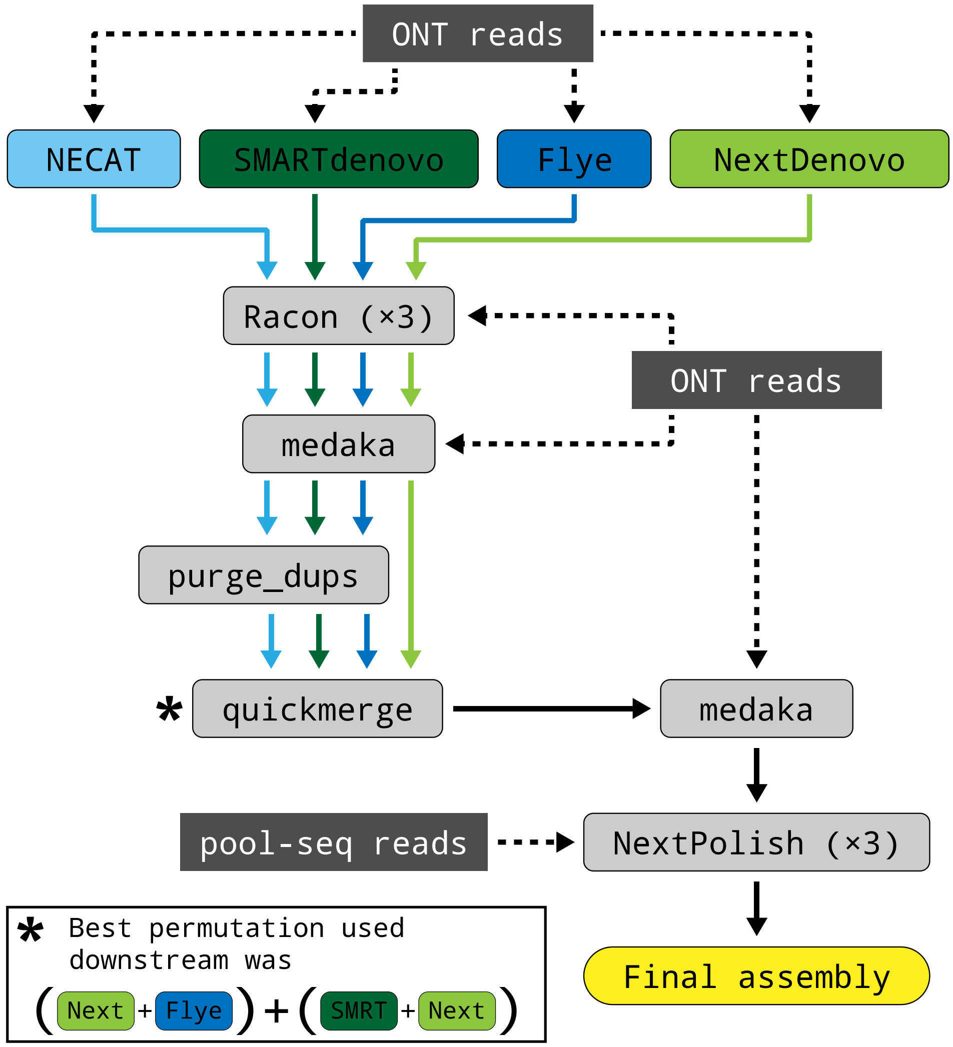

**Figure S5.** Pipeline to generate the *Tanytarsus gracilentus* reference assembly.

**Table S1.** Summary statistics for genome annotations. Included is completeness of coding sequences (CDS) calculated using BUSCO and the diptera_odb10 protein database. Also included are the number of genes with tags for Kyoto Encyclopedia of Genes and Genomes (KEGG) Orthology (KO) terms, Gene Ontology (GO) terms, Clusters of Orthologous Genes (COG) categories, Enzyme Commission (EC) numbers, and KEGG enzyme pathways.

| Feature | Assembly | | | |
|---|---|---|---|---|
| | *Chironomus riparius** | *Culicoides sonorensis* | *Parochlus steinenii* | *Tanytarsus gracilentus* |
| total proteins | – | 18,212 | 16,259 | 15,561 |
| protein-coding genes | 16,522 | 18,080 | 16,105 | 15,499 |
| % CDS complete | – | 90.0 | 95.1 | 93.4 |
| % CDS complete + single | – | 72.1 | 88.7 | 87.9 |
| % CDS complete + duplicated | – | 17.8 | 6.4 | 5.5 |
| KO terms | 10,576 | 10,760 | 9,675 | 9,753 |
| GO terms | 7,414 | 7,907 | 7,095 | 6,834 |
| COG categories | 3,937 | 4,411 | 4,009 | 3,785 |
| EC numbers | 3,698 | 3,870 | 3,517 | 3,566 |
| KEGG enzyme pathways | 3,080 | 3,427 | 3,074 | 2,927 |

* This species only had functional annotation created here so did not have CDS completeness scored

**Table S2.** Estimates and parametric-bootstrapped 95% confidence interval bounds for Pearson correlations between genome size and genome features computed using cor_phylo.

| | Feature | $r$ | $r_{2.5\%}$ | $r_{97.5\%}$ |
|---|---|---|---|---|
| | Protein-coding genes | 0.160 | −0.373 | 0.583 |
| | Intergenic content | 0.962 | 0.857 | 1.000 |
| | Mean intron length | 0.913 | 0.574 | 1.000 |
| Repeat elements | DNA | 0.842 | 0.526 | 0.975 |
| | LINE | 0.776 | 0.412 | 0.946 |
| | LTR | 0.814 | 0.482 | 0.956 |
| | SINE | 0.769 | 0.403 | 0.966 |
| | Unclassified | 0.873 | 0.541 | 0.997 |
| | Non-TE | 0.760 | 0.370 | 0.992 |

**Table S3.** List of the six HOG gene families that significantly expanded at the node separating Chironomidae from their most recent common ancestor in our phylogeny. Listed are the gene(s) for *Culex quinquefasciatus* and *Musca domestica* that fall within the HOG and the descriptions for these genes from each species' genome annotation.

| HOG | Species | Gene | Description |
|---|---|---|---|
| N0.HOG0000001 | *Culex quinquefasciatus* | CQUJHB008986 | cytochrome P450 6a14 |
| | *Culex quinquefasciatus* | CQUJHB009005 | cytochrome P450 6A1 |
| | *Culex quinquefasciatus* | CQUJHB009148 | cytochrome P450 6a14 |
| | *Musca domestica* | Mdom002048 | cytochrome P450 317a1 |
| N0.HOG0000121 | *Culex quinquefasciatus* | CQUJHB001249 | facilitated trehalose transporter Tret1-like |
| | *Culex quinquefasciatus* | CQUJHB005468 | facilitated trehalose transporter Tret1-like |
| | *Culex quinquefasciatus* | CQUJHB010329 | facilitated trehalose transporter Tret1 |
| | *Musca domestica* | Mdom004618 | Facilitated trehalose transporter Tret1-2 homolog |
| N0.HOG0000215 | *Culex quinquefasciatus* | CQUJHB000177 | serine protease gd |
| | *Culex quinquefasciatus* | CQUJHB000414 | serine protease gd-like |
| | *Culex quinquefasciatus* | CQUJHB001030 | CLIP domain-containing serine protease B15 |
| | *Culex quinquefasciatus* | CQUJHB004853 | coagulation factor IX-like |
| | *Culex quinquefasciatus* | CQUJHB005641 | serine protease gd |
| | *Culex quinquefasciatus* | CQUJHB008470 | chymotrypsin-like protease CTRL-1 |
| | *Culex quinquefasciatus* | CQUJHB013601 | transmembrane protease serine 12-like |
| | *Musca domestica* | Mdom003072 | Serine protease gd |
| N0.HOG0000153 | *Culex quinquefasciatus* | CQUJHB000433 | N-acetylgalactosaminyltransferase 6 |
| | *Culex quinquefasciatus* | CQUJHB009199 | N-acetylgalactosaminyltransferase 6 |
| | *Musca domestica* | Mdom000331 | N-acetylgalactosaminyltransferase 6 |
| | *Musca domestica* | Mdom003911 | N-acetylgalactosaminyltransferase 4 |
| | *Musca domestica* | Mdom011216 | N-acetylgalactosaminyltransferase 6 |
| | *Musca domestica* | Mdom012602 | N-acetylgalactosaminyltransferase 4 |
| N0.HOG0000323 | *Musca domestica* | Mdom013520 | ATP-binding cassette sub-family G member 1 |
| N0.HOG0000559 | *Culex quinquefasciatus* | CQUJHB009847 | glycoprotein hormone G-protein coupled receptor |
| | *Musca domestica* | Mdom011711 | Adenosine receptor A2a |

**Table S4.** Targeted Gene Ontology (GO) terms used in the analyses for positive selection.

| GO term | Description | Parent category |
|---|---|---|
| GO:0010038 | response to metal ion | response to stimulus ><br>response to chemical ><br>response to inorganic substance |
| GO:0010212 | response to ionizing radiation | response to stimulus ><br>response to abiotic stimulus ><br>response to radiation |
| GO:0034059 | response to anoxia | response to stimulus ><br>response to stress |
| GO:0009409 | response to cold | response to stimulus ><br>response to stress |
| GO:0009408 | response to heat | response to stimulus ><br>response to stress |
| GO:0001666 | response to hypoxia | response to stimulus ><br>response to stress |
| GO:0006979 | response to oxidative stress | response to stimulus ><br>response to stress |
| GO:0098542 | defense response to other organism | response to stimulus ><br>response to external stimulus ><br>response to stress ><br>response to other organism |

**Table S5.** Sources of all genome assemblies and annotations used in this study. Unless otherwise noted, each annotation source provides a set of both gene predictions and functional annotations with GO terms. Families are abbreviated as follows: Ch = Chironomidae, Ce = Ceratopogonidae, Cu = Culicidae, and Mu = Muscidae.

| | | Assembly | | Annotation | |
|---|---|---|---|---|---|
| Family | Species | Source | Accession | Source | Accession |
| Ch | *Tanytarsus gracilentus* | Present study | – | Present study | – |
| | *Chironomus riparius* | Rothamsted Research | GCA_917627325.3 | GenBank* | GCA_917627325.3 |
| | *Chironomus tentans* | (Kutsenko et al. 2014) | GCA_000786525.1 | InsectBase | IBG_00179 |
| | *Polypedilum vanderplanki* | (Yoshida et al. 2022) | GCA_018290095.1 | InsectBase | IBG_00656 |
| | *Polypedilum pembai* | (Shaikhutdinov et al. 2023) | GCA_014622435.1 | InsectBase | IBG_00655 |
| | *Belgica antarctica* | (Kelley et al. 2014) | GCA_000775305.1 | InsectBase | IBG_00108 |
| | *Clunio marinus* | (Kaiser et al. 2016) | GCA_900005825.1 | InsectBase | IBG_00191 |
| | *Propsilocerus akamusi* | (Sun et al. 2021) | GCA_018397935.1 | InsectBase | IBG_00659 |
| | *Parochlus steinenii* | Present study† | – | Present study | – |
| Ce | *Culicoides sonorensis* | (Morales-Hojas et al. 2018) | GCA_900258525.3 | Present study | – |
| Cu | *Anopheles stephensi* | (Chakraborty et al. 2021) | GCA_013141755.1 | VectorBase | GCF_013141755.1, Sep 08, 2020 |
| | *Aedes aegypti* | (Matthews et al. 2018) | GCA_002204515.1 | VectorBase | GCA_002204515.1, AaegL5.3 |
| | *Culex quinquefasciatus* | (Behura et al. 2011) | GCA_015732765.1 | VectorBase | GCA_015732765.1, Dec 04, 2020 |
| Mu | *Musca domestica* | (Scott et al. 2014) | GCA_000371365.1 | InsectBase | IBG_00558 |

* Functional annotation with GO terms was not found on GenBank, so it was generated in this study

† Based on reads from (Shin et al. 2019)

**Table S6.** Summary statistics for *Tanytarsus gracilentus* assemblies. In the “program” column, “>” indicates that this program was run on the output from the previous step, and the quickmerge assembly shown is the best permutation (see table S7).

| program | size (Mb) | # contigs | N50 (Mb) | min (bp) | max (Mb) | % BUSCOs (n = 3,285) |
|---|---|---|---|---|---|---|
| NECAT | 93.49 | 198 | 2.15 | 619 | 3.88 | C:88.65 [S:86.61, D:2.04], F:2.68, M:8.68 |
| > Racon | 93.60 | 192 | 2.16 | 1,062 | 3.90 | C:89.28 [S:87.28, D:2.01], F:2.62, M:8.10 |
| > medaka | 93.85 | 192 | 2.17 | 1,070 | 3.91 | C:90.20 [S:88.10, D:2.10], F:1.83, M:7.98 |
| > purge_dups | 91.90 | 121 | 2.13 | 2,004 | 3.91 | C:90.20 [S:88.68, D:1.52], F:1.80, M:8.01 |
| SmartDenovo | 95.11 | 159 | 1.40 | 7,167 | 5.34 | C:87.67 [S:85.18, D:2.50], F:3.53, M:8.80 |
| > Racon | 95.42 | 159 | 1.41 | 5,040 | 5.37 | C:89.77 [S:87.25, D:2.53], F:2.65, M:7.58 |
| > medaka | 95.47 | 159 | 1.41 | 5,077 | 5.37 | C:90.96 [S:88.22, D:2.74], F:1.77, M:7.28 |
| > purge_dups | 92.84 | 119 | 1.52 | 6,612 | 5.37 | C:90.87 [S:89.19, D:1.67], F:1.80, M:7.34 |
| NextDenovo | 93.03 | 88 | 2.23 | 16,933 | 6.77 | C:89.65 [S:88.46, D:1.19], F:2.34, M:8.01 |
| > Racon | 92.80 | 88 | 2.23 | 16,037 | 6.76 | C:89.80 [S:88.80, D:1.00], F:2.47, M:7.73 |
| > medaka | 92.61 | 88 | 2.23 | 16,049 | 6.76 | C:90.93 [S:89.95, D:0.97], F:1.74, M:7.34 |
| Flye | 97.11 | 1,114 | 0.93 | 495 | 3.70 | C:89.68 [S:87.06, D:2.62], F:2.44, M:7.88 |
| > Racon | 97.51 | 1,052 | 0.94 | 92 | 3.72 | C:89.71 [S:87.31, D:2.40], F:2.44, M:7.85 |
| > medaka | 97.87 | 1,052 | 0.94 | 92 | 3.74 | C:90.93 [S:88.49, D:2.44], F:1.80, M:7.28 |
| > purge_dups | 91.89 | 333 | 0.99 | 203 | 3.74 | C:90.93 [S:89.68, D:1.25], F:1.80, M:7.28 |
| quickmerge | 92.33 | 45 | 7.05 | 16,049 | 12.43 | C:91.05 [S:90.08, D:0.97], F:1.74, M:7.21 |
| > medaka | 92.03 | 45 | 7.02 | 16,040 | 12.40 | C:91.11 [S:90.20, D:0.91], F:1.74, M:7.15 |
| > NextPolish | 91.83 | 45 | 7.01 | 16,039 | 12.37 | C:91.60 [S:90.53, D:1.07], F:1.49, M:6.91 |

**Table S7.** Summary statistics for two rounds of quickmerge on initial *T. gracilentus* assemblies. The permutation column indicates the identity and order of assemblies input to quickmerge, and the lower set of permutations are the second round that use the four best first-round permutations.

| permutation | size (Mb) | # contigs | N50 (Mb) | min (bp) | max (Mb) | % BUSCOs |
|---|---|---|---|---|---|---|
| flye + necat | 91.45 | 180 | 6.05 | 203 | 13.96 | C:89.95 [S:88.65, D:1.31], F:1.80, M:8.25 |
| flye + next | 93.32 | 156 | 6.50 | 203 | 18.17 | C:90.41 [S:89.13, D:1.28], F:1.67, M:7.91 |
| flye + smart | 89.87 | 192 | 4.50 | 203 | 11.07 | C:88.19 [S:86.79, D:1.40], F:1.61, M:10.20 |
| necat + flye | 91.45 | 103 | 2.76 | 2,004 | 13.92 | C:90.11 [S:88.65, D:1.46], F:1.74, M:8.16 |
| necat + next | 93.77 | 106 | 2.77 | 2,004 | 6.76 | C:90.05 [S:86.70, D:3.35], F:1.77, M:8.19 |
| necat + smart | 92.41 | 96 | 3.07 | 2,004 | 13.30 | C:90.29 [S:88.25, D:2.04], F:1.70, M:8.01 |
| * next + flye | 92.34 | 68 | 3.31 | 16,049 | 12.43 | C:91.02 [S:90.05, D:0.97], F:1.74, M:7.25 |
| next + necat | 93.49 | 75 | 2.89 | 17,544 | 6.76 | C:90.08 [S:87.64, D:2.44], F:1.77, M:8.16 |
| * next + smart | 92.45 | 62 | 3.31 | 17,544 | 9.01 | C:90.96 [S:90.05, D:0.91], F:1.77, M:7.28 |
| smart + flye | 92.33 | 69 | 6.30 | 6,612 | 11.25 | C:90.87 [S:89.13, D:1.74], F:1.77, M:7.37 |
| * smart + necat | 92.47 | 53 | 5.31 | 6,612 | 13.24 | C:89.53 [S:87.00, D:2.53], F:1.74, M:8.74 |
| * smart + next | 92.11 | 56 | 5.38 | 6,612 | 11.63 | C:90.93 [S:89.92, D:1.00], F:1.70, M:7.37 |
| (next + flye) + (next + smart) | 94.80 | 60 | 6.05 | 16,049 | 12.43 | C:91.02 [S:87.61, D:3.41], F:1.74, M:7.25 |
| (next + smart) + (next + flye) | 92.29 | 54 | 6.05 | 17,544 | 12.43 | C:91.02 [S:90.08, D:0.94], F:1.74, M:7.25 |
| (next + flye) + (smart + necat) | 93.06 | 40 | 7.05 | 17,544 | 13.25 | C:90.62 [S:88.34, D:2.28], F:1.80, M:7.58 |
| (smart + necat) + (next + flye) | 90.21 | 41 | 8.58 | 6,612 | 14.25 | C:88.49 [S:86.88, D:1.61], F:1.70, M:9.80 |
| † (next + flye) + (smart + next) | 92.33 | 45 | 7.05 | 16,049 | 12.43 | C:91.05 [S:90.08, D:0.97], F:1.74, M:7.21 |
| (smart + next) + (next + flye) | 92.01 | 46 | 7.05 | 6,612 | 12.43 | C:90.96 [S:89.89, D:1.07], F:1.67, M:7.37 |

* first-round permutation used in second round
† best second-round permutation used as final quickmerge assembly

**Table S8.** Parameters defining the statistical distributions used to sample divergence times in MCMCTree, for calibrations defining (A) lower and upper and (B) just lower bounds. All values use 100 million years ago as the time unit because MCMCTree requires that all times be 0.01–10. (A) Parameter $t_L$ is the lower age bound, $t_U$ is the upper age bound, $p_L$ is the probability that the real age is less than $t_L$, and $p_U$ is the probability that the real age is greater than $t_U$. Upper and lower constraint specifications take the form 'B($t_L$, $t_U$, $p_L$, $p_U$)' in MCMCTree. (B) Parameter $p$ is the offset, $c$ is the scale parameter. Lower constraint commands take the form 'L($t_L$, $p$, $c$, $p_L$)'. In the Cauchy distribution used for sampling, the location parameter is $t_L[1 + p]$, and the scale parameter is $c\ t_L$. When a distribution mode was based on a reference, this means that we adjusted parameter $p$ so that the distribution mode matched the estimate in that reference.

| A. Lower and upper constraints: | | | | | |
|---|---|---|---|---|---|
| Node | $t_L$ | $t_U$ | $p_L$ | $p_U$ | Notes |
| a | 2.385 | 2.954 | 0.025 | 0.1 | Bounds based on (Benton et al. 2009), increased softness due to estimates in (Cranston et al. 2012) |
| B. Just lower constraints: | | | | | |
| Node | $t_L$ | $p$ | $c$ | $p_L$ | Notes |
| b | 2.420 | 0.1 | 0.2 | 0.1 | increased softness and a low distribution mean because of lower estimate in (Benton et al. 2009) |
| c | 2.013 | 0.16 | 0.5 | 0.025 | distribution mode based on (Cranston et al. 2012) |
| d | 0.935 | 0.47 | 0.5 | 0.025 | distribution mode based on (Cranston et al. 2012) |
| e | 0.339 | 1.24 | 1.0 | 0.025 | distribution mode based on (Benton et al. 2009) |
